# The genetic architecture of the sexually selected sword ornament and its evolution in hybrid populations

**DOI:** 10.1101/2020.07.23.218164

**Authors:** Daniel L. Powell, Cheyenne Payne, Mackenzie Keegan, Shreya M. Banerjee, Rongfeng Cui, Peter Andolfatto, Molly Schumer, Gil G. Rosenthal

## Abstract

Biologists since Darwin have been fascinated by the evolution of sexually selected ornaments, particularly those that reduce viability. Uncovering the genetic architecture of these traits is key to understanding how they evolve and are maintained. Here, we investigate the genetic architecture of a sexually selected ornament, the “sword” fin extension that characterizes many species of swordtail fish (*Xiphophorus*). Using sworded and swordless sister species of *Xiphophorus*, we generated a mapping population and show that the sword ornament is polygenic – with ancestry across the genome explaining substantial variation in the trait. After accounting for the impacts of genome-wide ancestry, we identify one major effect QTL that explains ∼5% of the overall variation in the trait. Using a series of approaches, we narrow this large QTL interval to a handful of likely candidate genes, including the gene *sp8*. Notably, *sp8* plays a regulatory role in fin regeneration and harbors several derived substitutions that are predicted to impact protein function in the species that has lost the sword ornament. Furthermore, we find evidence of selection on ancestry at *sp8* in four natural hybrid populations, consistent with selection against the sword in these populations.

## Introduction

The diversity generated by sexual selection poses an evolutionary puzzle. Why are courtship traits so different from one species to the next? Theoretical models suggest that much of the answer may hinge on the genetic architecture underlying sexual communication [1,2]. With the genomic revolution, we have made massive progress in understanding the genetic architecture of complex traits, particularly in humans. On the whole, this research has revealed that many traits, even those formerly assumed to have a simple genetic basis [3], are in fact highly polygenic, with hundreds to thousands of sites contributing to trait variation [4].

By contrast, we know far less about the genetic architecture of adaptive traits like sexual signals that arise over evolutionary timescales. Previous work has hinted at a simpler genetic basis for adaptive traits [5–7], including traits under sexual selection [8–10], however it is often challenging to disentangle variation in genetic architecture from variation in power to map traits of interest [11]. Moreover, statistical challenges like the winner’s curse [12] make it difficult to interpret the distribution of effect sizes in such studies.

Here, we investigate the genetic architecture and trace the evolutionary history of a well-studied sexually-selected trait in swordtail fish (*Xiphophorus*). The sword is a male-specific ornament generated by an elongation of the lower caudal fin rays (Fig 1). The sword ornament likely evolved in the last 3-5 million years [13,14] as a result of preexisting female mating preferences for the trait [15,16]. However, contemporary species vary widely in their expression of the ornament. The length of the sword ranges from complete absence to swords exceeding male body length (Fig 1; [17]). Female preference for swords also varies across species, from strong preference to antipathy towards swords [16,18], but is also impacted by social learning [19–21]. How the sword is predicted to evolve in response to female preferences depends in part on its underlying genetic architecture [2,22].

**Fig 1.**
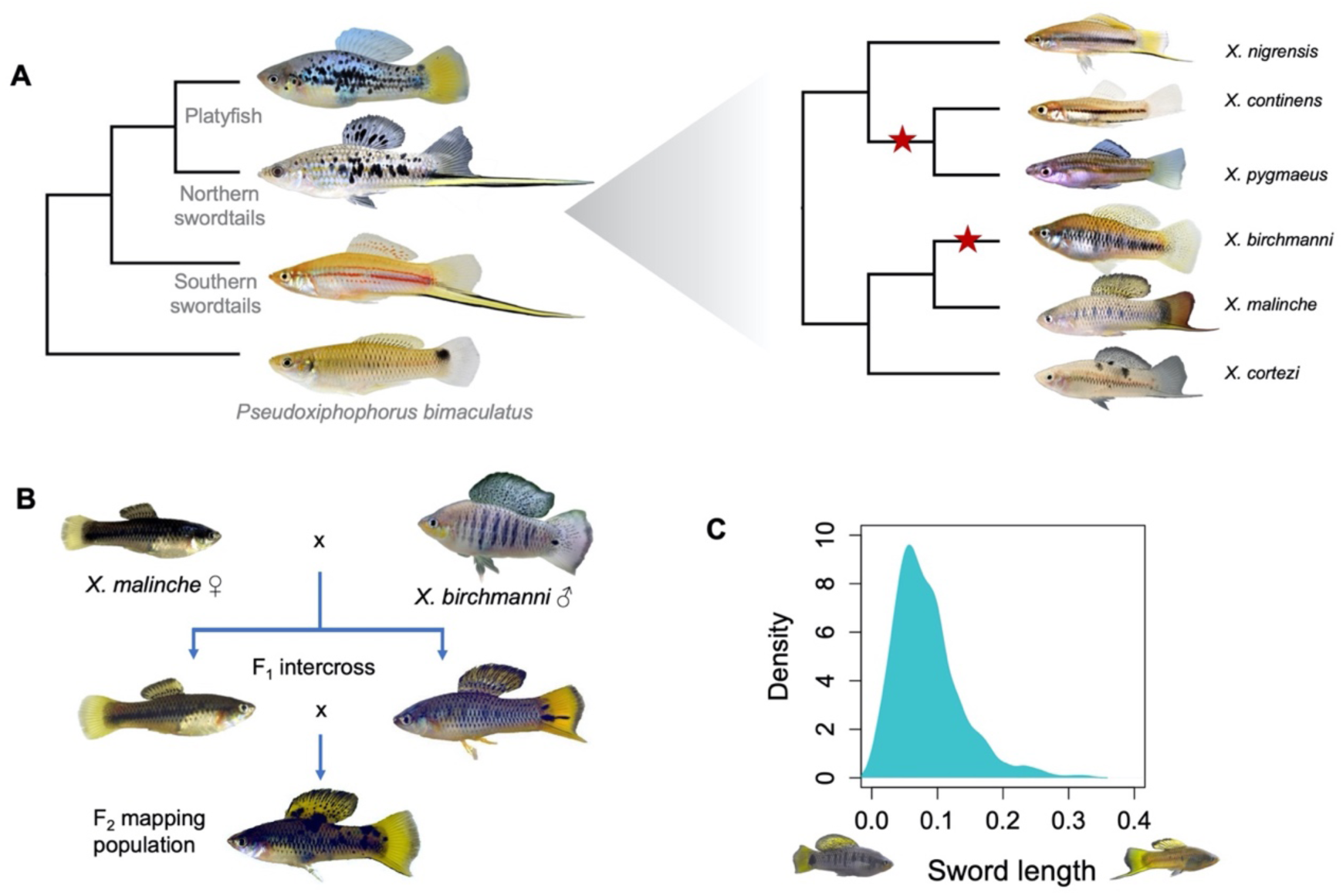
Evolutionary history of the sword and study design. **A**. Left - Phylogenetic relationships between platyfish, northern swordtails, and southern swordtails. The sword ornament was lost in the common ancestor of all platyfish. Right - Phylogenetic relationships among northern swordtails highlights at least two losses of the sword within this clade (red stars). **B**. Cross design used in this study involved crosses between *X. malinche* females and *X. birchmanni* males, followed by intercrosses between F_1_ hybrids. **C**. Distribution of normalized sword length in individuals within the hybrid mapping population. Photographs on the x-axis show an example of a hybrid individual with a normalized sword length of zero and an example of a hybrid individual with a normalized sword length of 0.35.

The complete loss of the sword in some *Xiphophorus* species affords the opportunity to characterize the genetic architecture of this sexually selected trait. In *Xiphophorus birchmanni*, males lack swords and females show strong preference for swordless males [23]. The absence of the sword in *X. birchmanni* is due to a recent loss of the trait, sometime after it diverged from its sister lineage, *X. malinche*, approximately 200,000 generations ago (or 100,000 years assuming two generations per year) [14,24]. In *X. malinche*, males have a pronounced sword ornament, but females paradoxically appear to prefer *X. birchmanni* visual phenotypes [20].

Like several pairs of species in the genus, *X. birchmanni* and *X. malinche* naturally hybridize [25,26]. Natural and artificial hybrid males vary in their sword phenotype, from swordless to swords as long as those of *X. malinche* (Fig 1). Given the importance of this novel trait in sexual selection theory, we sought to identify regions of the genome associated with the sword and understand their evolutionary history.

## Results

### Estimating the heritability of sword length in hybrids

Due to the fixed differences in sword phenotype between *X. birchmanni* and *X. malinche* and variable sword length in hybrids (Fig S1; mean sword length to body length ratio *birchmanni* = 0.016, *malinche* = 0.28), we knew that the sword was heritable. However, we wanted to quantify how much of the variation in sword length in hybrids could be attributed to genetic factors when individuals were raised in controlled conditions. To do so, we took advantage of a quantitative genetics based method for inferring the broad sense heritability of sword length by comparing phenotypic variance in F_1_ hybrids, where all individuals are genetically identical with respect to ancestry, to phenotypic variance in F_2_ hybrids [27] (Fig S1). We note that this approach assumes that all phenotypic variance in the parental species is due to environmental effects (see Materials & Methods for more details). This approach resulted in an estimate of 0.48 for broad sense heritability of the sword.

### Mapping the genetic basis of the sword phenotype

Although we have access to naturally occurring hybrids [28], we focused our mapping on artificial hybrids reared in common conditions (Materials & Methods), given that rearing condition can affect sword length [29]. We phenotyped 536 adult male F_2_ hybrids and collected low-coverage whole-genome sequence data (∼0.2X coverage; Materials & Methods). Using a pipeline we previously developed [30], we inferred local ancestry of each individual along the 24 swordtail chromosomes (Fig S2). Simulations indicated that we expect this approach to have high accuracy given our cross design (Fig S3; Supporting Information 1).

We thinned our initial dataset of 623,053 ancestry informative sites by physical distance to retain one marker per 50 kb for mapping. Ancestry linkage disequilibrium in lab-generated hybrids extends over several megabases. This resulted in 12,794 markers that were approximately evenly distributed along the genome (95% quantile of inter-marker distance, 60 kb; 98% of markers present in all individuals). Using the scanone function in R/qtl [31], we recovered one significant QTL for sword length on chromosome 13 (1.5 LOD interval = 4.1 Mb). As expected, individuals that harbored *X. malinche* ancestry in this region of chromosome 13 had longer swords on average, and the effects of the QTL appear to be additive (Fig 2). Bootstrapping and joint analysis with another study allowed us to narrow this large interval to 1.2 Mb ([32]; Supporting Information 2).

**Fig 2.**
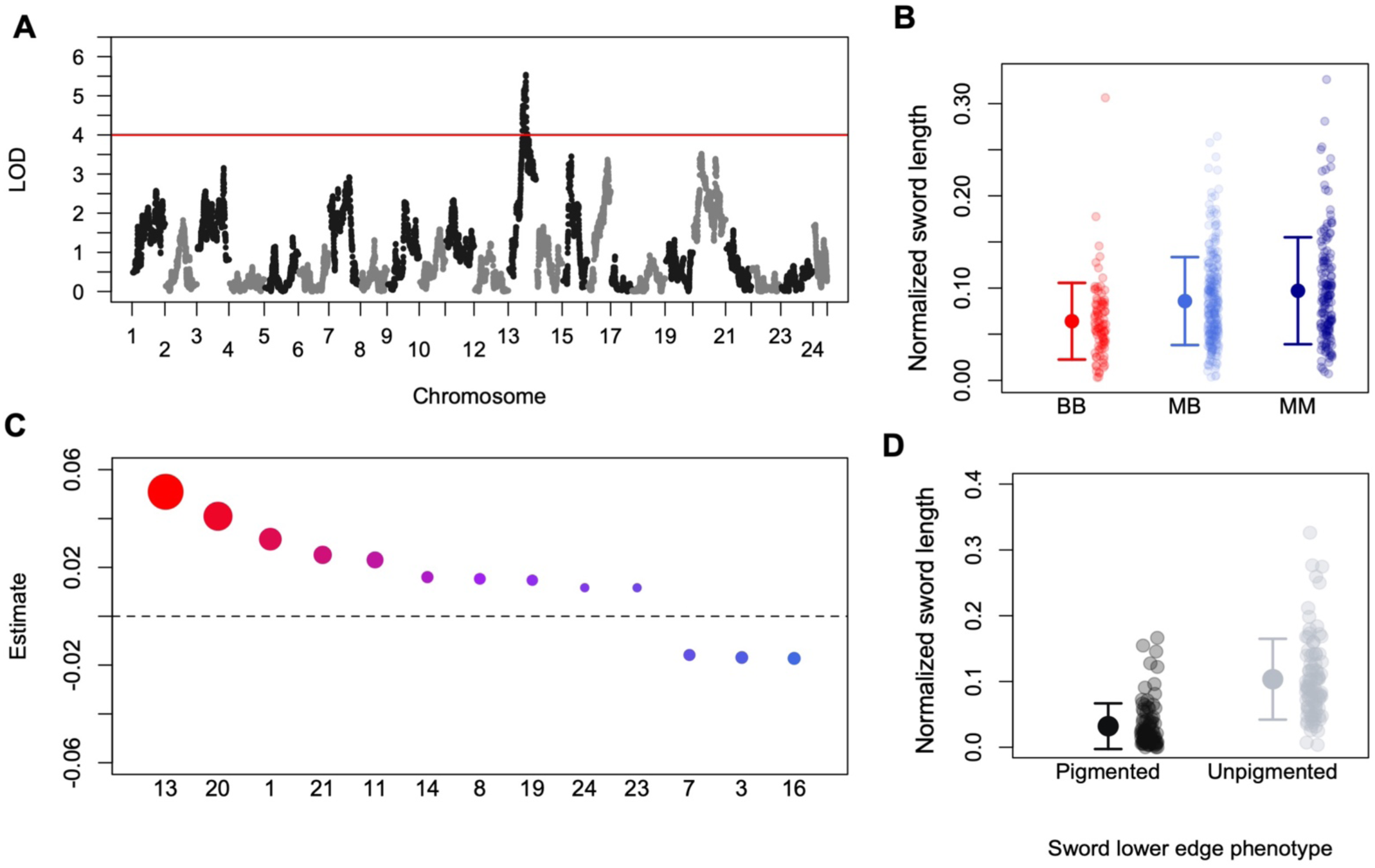
Ancestry at chromosome 13 and throughout the genome contributes to sword length. **A**. Manhattan plot of QTL mapping results for sword length reveal a single genome-wide significant QTL. Red line indicates genome-wide significant threshold determined by permutation; LOD – logarithm of odds. **B**. Sword length as a function of ancestry at the QTL peak. Small semi-transparent points show the raw data and large points and whiskers show the mean ± two standard errors of the mean. BB - homozygous *X. birchmanni*, MB - heterozygous, MM - homozygous *X. malinche*. **C**. Estimated effect sizes of ancestry on each chromosome using a model selection approach to select the minimal set of chromosomes that explain sword length. Point size corresponds to the p-value, with more significant associations represented as large points. **D**. The sword is a composite phenotype that includes a pigmented edge. Sword length is not strongly correlated with the presence of the lower edge pigmentation (Fig S4), suggesting an even more complex genetic basis of the composite trait than indicated by analyses of sword length alone.

Estimated effect sizes from QTL mapping studies are often inflated in cases where the experiment has low statistical power [12]. Aware of such issues, we used an approximate Bayesian computation (ABC) approach to estimate the range of effect sizes for the chromosome 13 QTL consistent with our data (Supporting Information 3). Based on this analysis, we estimate that *X. malinche* ancestry at the QTL peak on chromosome 13 explains 5.5% of the total variation in sword length (Fig S1; 95% confidence intervals: 1-22%) or approximately 11% of the heritable variation (Supporting Information 3). We found that the QTL region was syntenic between *X. birchmanni* and *X. malinche*, with no evidence for inversions, insertions, or deletions between the species (Fig S5).

Despite finding only a single significant genome-wide association with the sword, multiple lines of evidence indicate that the genetic architecture of the sword is more complex. Genome-wide ancestry is strongly associated with sword length (Spearman’s ρ = 0.2, p<4×10^−6^), and as expected individuals with a greater proportion of their genome derived from *X. malinche* tended to have longer swords (Fig S6). This association remains even after accounting for an individual’s ancestry within the QTL region of chromosome 13, indicating that it is not driven by the contribution of the QTL region to genome-wide ancestry variation (Spearman’s ρ = 0.18, p<4×10^−5^; Fig S6).

We next asked whether ancestry on particular chromosomes explained more of the variance in sword phenotype than ancestry on chromosome 13 or genome-wide ancestry (Materials & Methods). Based on regression analysis, we found that *X. malinche* ancestry on chromosomes 1, 11, 20 and 21 was significantly associated with variation in sword length, and together with chromosome 13 explained an estimated 27% of the heritable variation in the trait (Fig 2). We did not find a significant correlation between chromosome length and estimated effect size (R=-0.09, p=0.66). After accounting for ancestry on chromosomes 1, 11, 13, 20 and 21, *X. malinche* ancestry elsewhere in the genome was no longer significantly predictive of sword length in a partial correlation analysis. However, an AIC-based model selection approach retained thirteen chromosomes (54% of the genome) in the final model describing sword length as a function of chromosome level ancestry (Fig 2). QTL analysis including chromosome-level ancestry of each of the thirteen chromosomes retained in the AIC model as covariates yielded similar results (Fig S7). Surprisingly, although *X. malinche* ancestry was positively associated with sword length in most cases, *X. birchmanni* ancestry on three chromosomes was positively associated with sword length (Fig 2). Although these results suggest a lower bound for the number of regions underlying the sword phenotypes, approaches using the inferred effect size of the observed QTL indicate that the true number of causal loci could be much larger ([33]; Supporting Information 3).

Moreover, in addition to the genetic architecture of sword length, the sword itself is a composite trait [34]. The sword phenotype found in natural populations of *Xiphophorus* includes both the fin extension and a pigmented upper and lower margin (Fig 1, Fig 2). These traits can become decoupled in hybrids (Fig S4). Although sword length and upper sword edge are strongly correlated in hybrids (R=0.48, p<10^−32^), we detected a weaker correlation between the presence of the lower sword edge and sword length in hybrids (R= 0.19, p<10^−5^), even though both traits are always observed in sworded *X. malinche* (Supporting Information 4). Mapping attempts for the lower sword edge were unsuccessful. Specifically, no significant QTL peaks were identified, genome-wide ancestry was not strongly correlated with lower sword edge presence (R=0.05, p=0.25), and only ancestry on chromosome 11 was significantly associated with the lower sword edge (general linear model t=2.4, p=0.02). These results are discussed in more detail in Supporting Information 4.

### Functional data is consistent with several candidate genes within the chromosome 13 QTL

The sword normally develops during the course of sexual maturation in *X. malinche* and hybrid males. To evaluate evidence for possible candidates associated with sword length within the QTL region, we took advantage of the fact that adult male *X. malinche* will regenerate a complete sword if the sword tissue is removed (see Materials & Methods, Supporting Information 5, Fig S8). We also found that the sword will regrow in F_1_ hybrids, where all individuals have short swords (Fig S1, S8), but in *X. birchmanni* we simply observe regrowth of the normal caudal fin.

We reasoned that since the sword phenotype is recovered through the regrowth process [35,36], genes important in patterning and length should be expressed in the early stages of regrowth, and may be differentially expressed between *X. birchmanni, X. malinche*, and hybrids. Based on our RNA sequencing dataset (Materials & Methods, Supporting Information 5, Table S1), a large number of genes were differentially expressed between regenerating tissue in *X. birchmanni* and *X. malinche* (Fig 3; Fig S9). These differentially expressed genes were enriched for pathways including cell adhesion, cell cycle pathways, extracellular matrix-receptor interactions, and ribosome biogenesis (Materials & Methods, Supporting Information 3, 5). The first three pathways encompass a substantial number of genes with significant expression differences (Materials & Methods, Supporting Information 5). Many of the expression differences we observe during regeneration appear to be consistent with evolved changes in expression (i.e. allele-specific expression, Supporting Information 5).

**Fig 3.**
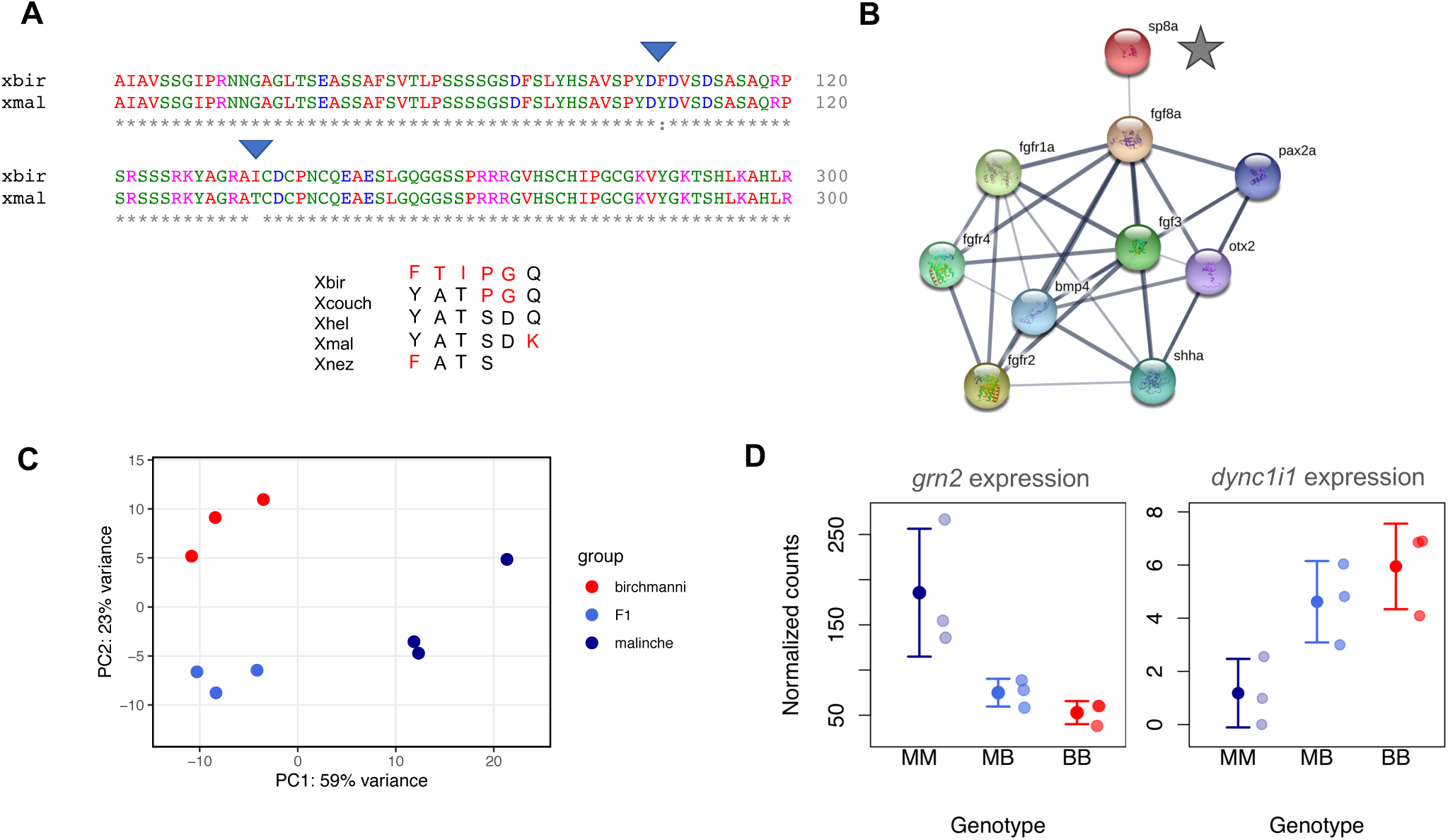
Expression and substitution data highlight candidate genes likely to drive variation in sword length. **A**. Clustal alignment of a section of the *X. birchmanni* and *X. malinche sp8*, which is found within the chromosome 13 QTL peak. Derived substitutions in *X. birchmanni* that are predicted to not be tolerated in a SIFT analysis are indicated by the blue triangles. Asterisks indicate identical amino acid sequences, colors indicate amino acid properties, blanks, colons and periods indicate substitutions. Below the alignment is a table of amino acid state at all sites that are variable in at least one of the species analyzed. Black indicates amino acids that follow the inferred ancestral state and red indicate amino acids that are derived. **B**. STRING network for the ortholog of *Xiphophorus sp8* in zebrafish (*sp8a*) shows that this gene regulates a number of fibroblast growth factor and fibroblast growth factor receptor genes (*fgf* and *fgfr* genes). These genes have been implicated in fin growth in zebrafish [37], limb development in other species [38], and were previously identified as likely candidates underlying sword regeneration in southern swordtails [35]. **C**. Principal component analysis of RNAseq data from regenerating caudal fin tissue, which will become sword tissue in *X. malinche* and F_1_s. **D**. Expression patterns of two candidate genes within the QTL region that are differentially expressed between *X. birchmanni* and *X. malinche* in the regenerating sword and show expression patterns consistent with predicted phenotypes (Table S3), although we observe low overall expression for *dync1i1*. BB - *X. birchmanni*, MB - F_1_ hybrids, MM - *X. malinche*. Small semi-transparent points show the raw data and solid points and whiskers show the mean ± two standard errors of the mean. Note that *sp8* is expressed but not significantly differentially expressed between species in regenerating caudal tissue (Fig S10).

Although there is evidence from ancestry and expression data that the production of the sword involves genetic differences on many chromosomes and substantial differences in expression response between species, we were interested in narrowing down likely candidates associated with sword length within the QTL region of chromosome 13. Of 52 genes in this region (Table S2), 16 had annotations associated with growth, skeletal, muscle, or limb development phenotypes. We further evaluated these candidates using a combination of differential expression and sequence analysis approaches (Supporting Information 3, 5), ultimately narrowing to eight genes most likely to drive variation in sword length due to their expression or substitution patterns (Table S3, Fig 3).

Of these genes, we highlight three of the most compelling candidates. *sp8*, which impacts limb and fin differentiation [37,38], has five derived nonsynonymous substitutions in *X. birchmanni* (Fig 3) and an overall rapid rate of protein evolution between *X. birchmanni* and *X. malinche* (*dN/dS* = 0.78; upper 3% genome-wide) [32]. Although *sp8* harbors a large number of substitutions derived in *X. birchmanni* (Fig 3), we did not find strong evidence for a different substitution rate along the *X. birchmanni* branch based on PAML analysis (χ^2^=3, p=0.08). However, two of the substitutions derived in *X. birchmanni* are predicted to affect protein function based on analysis with the program SIFT (Fig 3, Materials & Methods, [39]). We also evaluated these metrics for the other candidate genes in the region with amino acid substitutions between *X. birchmanni* and *X. malinche* (Table S4).

Two other genes, *dync1i1* and *grn2*, are differentially expressed in regenerating caudal tissue, and their expression patterns mirror predicted phenotypic differences between *X. birchmanni* and *X. malinche* (Fig 3). Misexpression of *dync1i1*, which is downregulated in regenerating sword tissue in *X. malinche*, is associated with abnormal limb and fin growth in other species ([40]; Fig 3). *grn2* is strongly upregulated in regenerating *X. malinche* fin tissue (Fig 3) and belongs to a family of progranulin growth factors which are implicated in regulating cell growth, proliferation, and regeneration [41].

### Sword QTL in hybrid populations

Given the importance of the sword as a sexually selected signal, we predicted that regions underlying variation in this phenotype may have unusual patterns of ancestry in natural hybrid populations formed between *X. birchmanni* and *X. malinche*. Behavioral research has indicated that in addition to males having lost the sword phenotype, female *X. birchmanni* prefer swordless males [23]. Although *X. malinche* males are sworded, females of this species appear indifferent to the sword and generally prefer *X. birchmanni* visual phenotypes [20,42]. Thus, we may expect that genomic regions underlying the sword would be selected against in hybrid populations, if swordless males, on average, have a mating advantage over sworded males.

We examined local ancestry around the chromosome 13 QTL in four hybrid populations using a combination of previously collected data and new data [24,43]. Two of these populations (Acuapa and Aguazarca) derive the majority of their genomes from the *X. birchmanni* parental species and two populations (Tlatemaco and Chahuaco falls) derive the majority of their genomes from the *X. malinche* parental species. Newly collected data for the Acuapa population is available through the NCBI sequence read archive (SRAXXXXXX, pending).

Overall, *X. birchmanni* x *X. malinche* hybrid populations do not show unusual ancestry in the chromosome 13 sword QTL region as a whole (Fig S11). However, given the size of the QTL, there is substantial heterogeneity in ancestry within the QTL region in natural hybrid populations where ancestry linkage disequilibrium decays over ∼1 Mb [24,43]. Interestingly, *X. malinche* ancestry is lower than expected across four independent hybrid populations around the gene *sp8* and those closely linked to it (p=0.002 by simulation, Materials & Methods, Table S4). This is notable because *sp8* was identified as a promising candidate within the chromosome 13 QTL region based on its phenotypic effects on fin and limb growth and the presence of amino acid substitutions likely to impact protein function between *X. birchmanni* and *X. malinche* (Fig 3; Table S3; Supporting Information 3). If *sp8* is indeed the causal locus within the QTL region, low *X. malinche* ancestry could be consistent with selection against the sword in hybrid populations.

We combined local ancestry analyses in population samples with another source of data that could also highlight selection in this region. For one of the hybrid populations studied above (Acuapa) we were able to develop a time-transect dataset, spanning an estimated 24 generations of hybrid population evolution (from 2006 to 2018). Although we do not see evidence for unusual changes in ancestry in the QTL region as a whole, we observe a decline in *X. malinche* ancestry over time within the Acuapa population at *sp8* (Fig 4), consistent with moderate selection against *X. malinche* ancestry in this region (maximum a posteriori estimate: −0.1, 95% confidence intervals: −0.44 to −0.03; Fig 4; Supporting Information 6). This direction of change in ancestry is opposite what would be expected due to population demography, given that the Acuapa population receives *X. malinche* but not *X. birchmanni* migrants ([28]; see Supporting Information 6). Other candidate genes in this region do not change significantly in ancestry over this time period, apart from genes with the strongest physical linkage to *sp8* (*sp4*, 11 kb away and *twistnb*, 60 kb away; Table S5).

**Fig 4.**
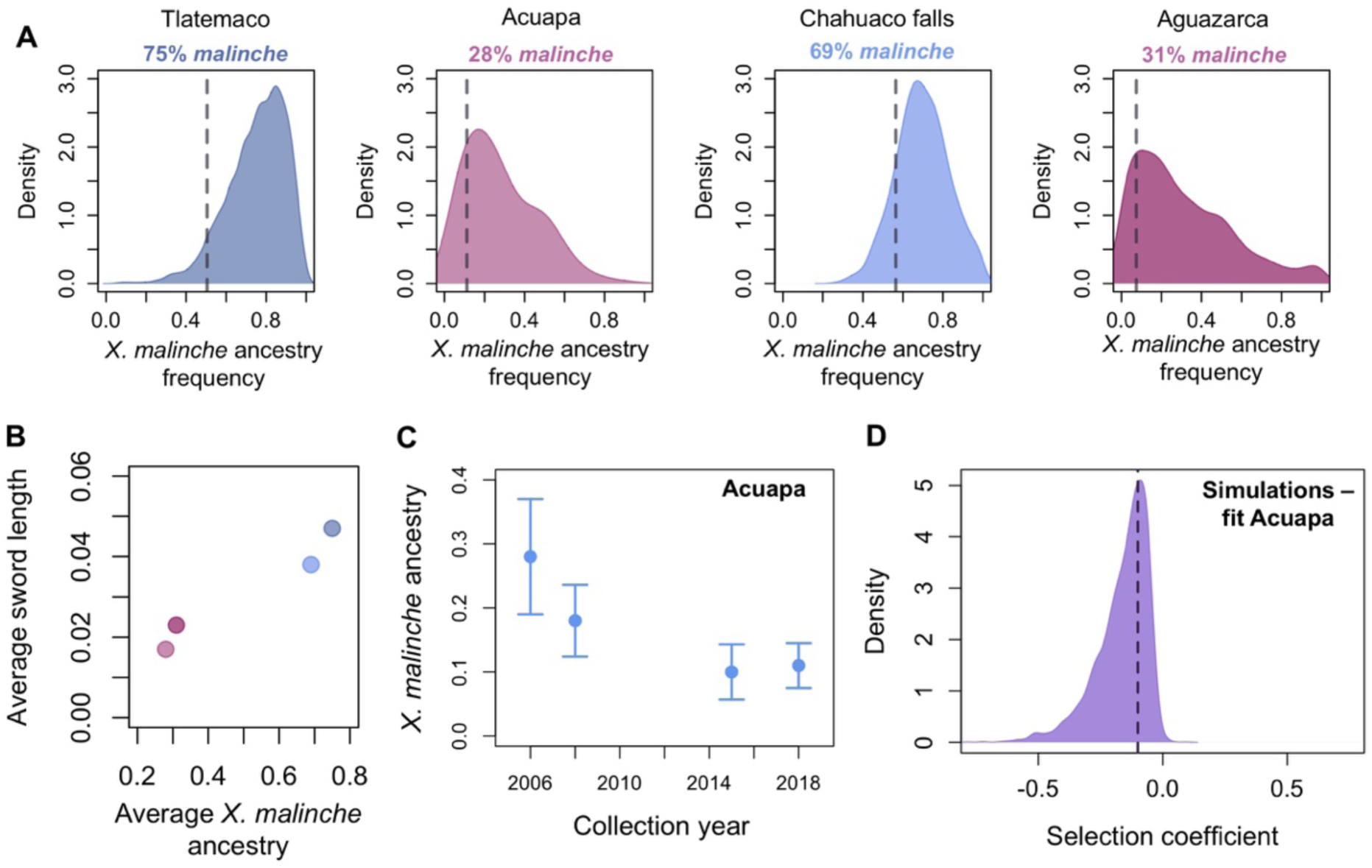
Patterns of ancestry at *sp8* are consistent with selection against *X. malinche* ancestry. **A**. Genome wide ancestry distribution in naturally occurring hybrid populations versus ancestry at *sp8* (dotted line). Across hybrid populations, ancestry at *sp8* falls in the lower tail of the *X. malinche* ancestry distribution. Ancestry is summarized genome wide and at *sp8* in 50 kb windows. **B**. Overall, observed sword length correlates with genome-wide ancestry across hybrid populations (Pearson’s R=0.98, p=0.03). Plotted phenotypes are based on 48-193 individuals per population. We caution that this analysis does not control for potential differences in environmental effects across populations. **C**. *X. malinche* ancestry decreases over time at *sp8* in a hybrid population where time series data is available (the Acuapa population). **D**. This decrease is consistent with moderate to strong selection against *X. malinche* ancestry at the *sp8* region in this population. Shown here is the posterior distribution of accepted parameters from ABC simulations (Supporting Information 6). Dashed line shows the maximum a posteriori estimate.

### Evolutionary patterns associated with the sword QTL

The distribution of the sword trait among *Xiphophorus* species indicates that there have been multiple losses of the trait [13,14]. Most species lacking a sword fall within the platyfish clade, representing an ancient loss of the trait (Fig 1). By contrast, the loss of the sword in *X. birchmanni* occurred since its divergence with *X. malinche*, an estimated 200,000 generations ago [24].

Given the distinct timescales and independence of these losses, we were surprised to find that a sword QTL on chromosome 13 is also identified using crosses between the southern swordtail species *X. hellerii* and the platyfish species *X. maculatus*, largely overlapping with our signal (Supporting Information 2). Because of extensive hybridization in the group, this led us to ask whether there was evidence of introgression of genes associated with the absence of the sword.

The ranges of *X. birchmanni* and *X. birchmanni* x *X. malinche* hybrids overlap with a single platyfish species, *X. variatus* [17,25]. Like other platyfish, *X. variatus* lacks the sword. We recently detected evidence of introgression from the lineage leading to *X. variatus* into *X. birchmanni* and *X. malinche* [24]. We asked whether *X. birchmanni* harbored platyfish derived ancestry tracts that coincided with the chromosome 13 sword QTL, and were not found in *X. malinche*. We used the program PhyloNet-HMM to identify such regions [44]. Based on simulations, we predict that this approach will have good power to detect fixed ancestry tracts, likely due to the large sequence divergence between the groups (Supporting Information 7). Notably, we do not detect any such tracts in the QTL peak near *sp8* or unusual phylogenetic relationships in this region (Fig S12). We confirmed this result with an F_4_ ratio-based approach which may be more robust to short ancestry tracts ([45], Supporting Information 7).

Together, this implies that introgression from *X. variatus* at the chromosome 13 QTL is not responsible for the loss of the sword in *X. birchmanni*. We caution, however, that we have not excluded a role for introgression at other, as of yet unknown regions, as a cause of recent sword loss in the *X. birchmanni* lineage.

## Discussion

Using a combination of approaches, we identified a major effect locus contributing to phenotypic variation in the length of the sword, a sexually selected trait that evolved in the common ancestor of *Xiphophorus* fish. First highlighted by Darwin, this trait has long served as a classic example in sexual selection theory of the role of female preferences in driving the evolution of male ornamentation [46]. Among eight plausible candidates in this region (Table S3), we highlight *sp8*, which is expressed in regenerating swords and is surprisingly divergent in sequence between *X. birchmanni* and *X. malinche* (Fig 3, Fig S10). In vertebrates, knockouts of *sp8* have truncated limb phenotypes [38,47,48]. Moreover, *sp8* regulates members of the fibroblast growth factor (*fgf*) signaling pathway ([38]; Fig 3), which has previously been implicated in fin growth in general [37,49] and sword growth in particular [35].

In addition to chromosome 13, we find that ancestry throughout the rest of the genome contributes to variation in sword length. Model selection suggests that sword length is explained by ancestry proportions on as many as 13 of 24 chromosomes (Fig 2). This includes the putative sex chromosome, although the estimated effect size is small (<1% of the variation explained; Fig 2). Moreover, during sword regrowth a suite of genes are differentially regulated (Fig 3) and some of this response is likely attributable to evolved expression differences between species (Supporting Information 5). Further, sword length is just one of several traits contributing to the composite sword ornament, including melanocyte pigmentation of the sword edge, and these traits can become decoupled in hybrids (Fig 1; Supporting Information 4). Together, these results highlight the complex genetic architecture of the sword phenotype.

Surprisingly, we also observe that *X. birchmanni* ancestry on several chromosomes is positively correlated with sword length. This result is puzzling since *X. birchmanni* males lack a sword and thus *X. birchmanni* ancestry should not contribute to longer swords in hybrids. Simulations suggest that these results are not expected to be an artifact of our analysis approach (Supporting Information 8). Instead we speculate that they could be explained by the predictions of Fisher’s geometric model [50,51], where different phenotypic optima in the two species (i.e. sworded and swordless males) result in fixation of different suites of genetic variants, whose combinatorial effects are uncovered in hybrids [52]. These observations highlight a general problem for QTL mapping approaches using interspecific hybrids, where the phenotypic variance observed in hybrids is not necessarily generated by the same set of loci responsible for trait differences between the parental populations. Indeed, given the frequency of transgressive traits in hybrids, such effects may be relatively common [26,53].

Our results here also serve to underscore an important finding from previous work that has been largely overlooked in the recent mapping literature [54]. Without accounting for variation in genome-wide ancestry in hybrids, we originally detected three genome-wide significant QTLs (Fig S13, Supporting Information 8). Examination of two of these associations, on chromosomes 1 and 20, revealed relatively flat peaks spanning most of the chromosome (Fig S13). After accounting for genome-wide ancestry, both signals fell below our genome-wide significance threshold (Fig 2, Fig S13, Supporting Information 8). Our simulations suggest that when traits are polygenic and there is realistic variation in ancestry among artificial hybrids, ignoring genome-wide ancestry can result in inflated QTL peaks (Supporting Information 8). This phenomenon was explored in earlier theoretical work from Visscher & Haley [54]. The underlying biological issue is that although individuals generated by artificial crosses have a certain proportion of their genome derived from each parental species (i.e. 50% in our study, Fig S14), substantial variation in genome-wide ancestry is generated by recombination. The technical issues that arise from this variance are analogous to those long-appreciated in the admixture mapping literature [55]. Here, we emphasize again this important issue and how overlooking it can potentially lead to misinterpretation of mapping results.

The loss of the sword in *X. birchmanni* is one of several losses in the genus (Fig 1) and there is an overall trend towards preference for reduced sword length in the genus as a whole [14,56]. Given the history of gene flow in *Xiphophorus*, even between distantly related species, we asked whether there was evidence of gene flow into *X. birchmanni* at the chromosome 13 QTL from a swordless species. Our results indicated that introgression from swordless species on chromosome 13 is unlikely to explain the loss of the sword (Fig S12; Supporting Information 7). We note again, however, that given the polygenic basis of the sword we cannot exclude a role of introgression in sword loss at other regions of the genome.

Results from natural populations suggest that there may be selection against the sword in hybrid zones formed between *X. birchmanni* and *X. malinche*. Female *X. birchmanni* prefer swordless males and female *X. malinche* appear to be indifferent to the sword [20,23]. Moreover, natural selection is expected to act in concert against sworded males, as they are more visually obvious to potential predators [57]. This leads to the expectation that there may be selection against *X. malinche* ancestry in regions associated with the sword. Interestingly, one of the candidate genes we identified, *sp8*, has lower than expected *X. malinche* ancestry across hybrid populations and decreases in *X. malinche* ancestry over time in one hybrid population (Fig 4; Table S5; Supporting Information 6). This decrease is consistent with moderate selection against *X. malinche* ancestry in this region (Fig 4). However, we caution that ancestry at the chromosome 13 QTL explains only a fraction of the overall trait variation; ancestry changes at other underlying loci may support different patterns of selection and trait evolution in hybrid populations.

The causes of variability in sexual ornamentation within and between species remains a source of controversy. Theory and empirical evidence suggest a modest role for so-called “good genes” selection, where ornaments predict offspring success [58]. By contrast, ornaments can rapidly evolve simply because they are attractive, if they exploit a preexisting bias or coevolve with female preferences [59,60]. While the predictions of the good genes model are not strongly dependent on genetic architecture, the dynamics of coevolutionary models depend critically on the underlying genetic architecture [60]. For example, theory predicts that only traits with a polygenic basis are likely to be driven to extreme exaggeration through so-called “runaway” sexual selection [1,2,22]. To date, however, most genetic studies of sexual ornaments have identified single loci of large effect on sex chromosomes [8–10].

Our results support a polygenic, largely autosomal, architecture underlying variation in the sword ornament, with contributions from ancestry on thirteen chromosomes. This polygenic genetic basis is consistent with a number of evolutionary mechanisms that have been proposed to explain the evolution of extraordinary diversity in sword phenotypes across *Xiphophorus*. Our findings contrast with some previous observations that identified a simpler genetic architecture for several sexually selected ornaments [8–10], highlighting the challenges ahead in understanding the genetic basis of many evolutionarily important traits.

## Materials & Methods

### Artificial crosses between *X. birchmanni* and *X. malinche*

We crossed *X. malinche* (female) x *X. birchmanni* (male) to produce an F_1_ generation; previous attempts to produce viable offspring from the reciprocal cross were largely unsuccessful. We reared virgin *X. malinche* (n = 24) born to females collected at the Chicayotla locality on the Río Xontla using baited minnow traps. Wild *X. birchmanni* sires (n = 10) were collected from the Coacuilco locality on the Río Coacuilco. The resulting F_1_ offspring from this cross were reared to maturity. Based on past experience, we knew that it would be difficult to generate large numbers of early-generation hybrids in the lab. As a result, in June of 2016 we seeded each of 29 mesocosm tanks with F_1_ hybrids (n = 21 per tank). These mesocosm tanks are 2000 L outdoor tanks kept in semi-natural conditions but protected from predators and fed once daily.

We sampled the mesocosms in January and May of both 2017 and 2018, at which time all adult males were anesthetized with tricaine methanesulfonate (Texas A&M IACUC protocol #2016-0190), marked individually with color-coded elastomer tags for future identification (Northwest Marine Technologies), photographed for phenotyping, and fin-clipped for genotyping before returning them to mesocosm tanks. In total we genotyped and phenotyped 536 adult early generation hybrid males. Analysis of crossover numbers indicate that the majority of these individuals are F_2_ hybrids, and we found that our results are robust to excluding likely later generation hybrids (Supporting Information 9).

### Phenotyping approaches

We measured standard length (distance from the tip of the mandible to the midpoint of the distal edge of the caudal peduncle), sword extension (distance from the edge of the caudal fin to the tip of the sword) [61] from photographs of adult males using the ImageJ software package [62]. For analysis, sword extension was standardized by dividing by standard length and is referred to as sword length throughout the manuscript. We note that sword length is usually referred to the distance from the caudal fin base to the sword tip, which differs from our terminology here for convenience.

### Low coverage whole genome sequencing

We used the Agencourt bead-based protocol (Beckman Coulter, Brea, California) to extract DNA from fin clips. We followed the manufacturer’s instructions for the extractions except that we used half reactions. DNA was diluted to 10 ng/µl and 5 µl of sample was mixed with Tn5 transposase enzyme pre-charged with adapters. This mixture was incubated at 55 °C for 7 minutes to enzymatically shear DNA and the reaction was stopped by adding 2.5 µl of 0.2% SDS and incubating at 55 °C for another 7 minutes. Three microliters of each sample were combined with a plate-level i5 index and one of 96 i7 indices in an individual PCR reaction using OneTaq HS Quick Load mastermix. After amplification, 5 µl of each library were pooled and the pool was purified using Agencourt AMPpure XP purification beads. Libraries were quantified using a Qubit fluorimeter (Thermo Scientific, Wilmington, DE). Libraries were evaluated for size distribution and quality using a Bioanalyzer 1000 (Agilent, Santa Clara, California). Libraries were sequenced on the Illumina HiSeq 4000 at Weill Cornell Medical Center across three lanes to collect paired-end 100 nucleotide reads. This data has been deposited on the NCBI sequence read archive (SRAXXXXXX, pending).

### Local ancestry inference

To infer local ancestry, we used a pipeline we previously developed called *ancestryinfer* [30,63]. Briefly, for each individual Illumina reads were mapped to both the *X. birchmanni* and *X. malinche* reference genomes; uniquely mapping reads were retained and counts for each allele were tabulated at each ancestry informative site. A hidden Markov model [63] was applied to these counts to generate posterior probabilities of each ancestry state (homozygous *birchmanni*, heterozygous, and homozygous *malinche*) at ancestry informative sites throughout the genome. This resulted in posterior probabilities at 623,053 sites genome-wide in our dataset.

For downstream analyses, we converted these posterior probabilities to hard calls. If an individual had a posterior probability greater than 0.9 for any ancestry state, they were assigned that ancestry state at the focal marker. On average artificial hybrids derived 50% of their genomes from each parental species, as expected from the cross design (Fig S14). Local ancestry also mirrored expected patterns for early generation hybrids (Fig S2). Simulation results suggest we expect to have high accuracy in local ancestry inference (Fig S3).

### Estimates of heritability

To estimate the broad sense heritability of the sword length trait, we took advantage of phenotypic data from F_1_ and F_2_ hybrids raised in common conditions [64]. We calculated the variance in normalized sword length contributed by environmental factors (V_E_) as the trait variance in F_1_ hybrids, where all individuals have identical ancestry states throughout the genome. We calculated the combined impacts of environment and genetic variance (V_G_) using phenotypic variance (V_P_) in F_2_ hybrids. This allowed us to solve for V_G_ and estimate broad sense heritability using the relationship *h*^2^_broad_ = V_G_/ V_P_ (see [64]).

We note that the approach that we use to estimate heritability was designed for inbred lines and assumes that phenotypic variation within the parental species is due to environmental variation. While this is likely a valid assumption for *X. birchmanni* (mean sword length normalized by body length = 0.016 ± 0.02), it may not be the case in *X malinche* where we observe greater variation in sword length (mean sword length normalized by body length = 0.28 ± 0.07). Thus, we evaluated possible impacts of genetic variation for sword length within *X. malinche* on heritability inference using simulations (Supporting Information 10). These simulations suggest that additional genetic variation for sword length within *X. malinche* is unlikely to strongly bias our estimates of the heritability driven by *X. malinche* ancestry (Supporting Information 10, Fig S15).

### QTL analysis

For QTL analysis, the data were thinned to retain one marker per 50 kb; this resulted in 12,794 markers spread approximately evenly across the genome (95% of intermarker distances were less than 60 kb). This thinning is necessary due to the computational intensity of analysis using the R/qtl software. We used the scanone function of R/qtl to perform single QTL model standard interval mapping using the EM algorithm [31]. Recombination fraction was estimated using the est.rf() function and markers missing genotype data were excluded using the drop.nullmarker() function. Since genome-wide hybrid index was significantly correlated with sword length (ρ = 0.20, p < 4×10^−6^) we included it as a covariate during mapping (see also Fig S13; Supporting Information 8). We also repeated mapping analysis including ancestry on each chromosome retained in AIC model selection as a covariate (Fig S7). Rearing tank and tank location were omitted as covariates because they did not affect phenotype distribution. The threshold for genome-wide logarithm of odds (LOD) at a false discovery rate of 5% was determined based on 1,000 permutations of sword phenotype onto observed genotypes. For the identified QTL, the region that fell within 1.5 LOD of the peak LOD value was treated as the associated interval for downstream analyses.

For each chromosome containing a significant QTL, we aligned that chromosome from the *X. birchmanni* and *X. malinche* assemblies [43] using the program MUMmer [65]. We found no evidence of structural rearrangements or deletions between the two species in this region (Fig S5).

### Genetic architecture of the sword

In addition to QTL mapping, we asked about genome-wide associations between sword length and ancestry. We summarized ancestry per chromosome and genome-wide and used a partial correlation approach with the ppcor method in R to identify chromosomes in which ancestry was significantly associated with sword length, after accounting for ancestry on chromosome 13. We adjusted p-values with a bonferroni correction for the number of chromosomes. Finally, we evaluated associations between ancestry and sword length using an AIC model selection approach. We input a model in which ancestry on all chromosomes was included as independent variables and used the *step* function in R to select the minimal model of sword length as a function of chromosome level ancestry.

We also evaluated whether features such as chromosome length, number of genes per chromosome, and number of differentially expressed genes per chromosome correlated with the estimated effect sizes of the 24 chromosomes (see also Supporting Information 5). We did not see a correlation between the number of annotated genes per chromosome and the estimated effect size of that chromosome, whether we included all chromosomes (Spearman’s ρ = 0.1, p = 0.6) or only those retained during model selection (Spearman’s ρ=0.57, p = 0.1), although the trend observed for the latter is suggestive.

### Sword regeneration experiments

In order to compare gene expression patterns in developing caudal tissue of *X. birchmanni, X. malinche*, and their F_1_ hybrids, we took tissue samples after ten days of regeneration from three pools of ten individual males for each genotype class (90 fish in total) following Offen et al. [35]. Samples had to be pooled due to the expectation of low RNA yield from individual samples [Manfred Schartl, personal communication]. Briefly, to begin the experiment we anesthetized each fish in MS-222 and amputated the distal 1/4 of the caudal fin, which includes the sword in *X. malinche* and F_1_ hybrids, using a sterile razorblade. After recovery from anesthesia, each fish was housed individually in 11-liter aquaria and fins were allowed to regenerate for ten days at 22°C. After ten days, each fish was once again anesthetized and the regenerating blastema was removed. The dissected tissue was divided into three sections, with the most ventral section corresponding to regenerated sword tissue in *X. malinche* and F_1_ hybrids. The ventral tissue sections were then pooled in groups of ten according to genotype and replicate for RNA extraction. We generated a total of three pools (30 males) for each biological condition: *X. birchmanni, X. malinche* and F_1_ hybrids. RNA was extracted from the pooled tissue using a Trizol based protocol followed by on-column DNAse treatment and purification using the Qiagen RNeasy Mini Kit (Qiagen, Valencia, CA). RNAseq libraries were prepared in a single batch by the Bauer Core at Harvard using the KAPA mRNA HyperPrep Kit (Roche, Palo Alto, CA) with 300-500 nanograms of input RNA. Samples were sequenced across two HiSeq 2000 lanes at Harvard’s Bauer Core (Table S6) and yielded 150bp paired-end reads.

### Differential expression analysis

We tested for differential expression between *X. birchmanni* and *X. malinche* in the libraries described above. We used the Cutadapt and FastQC wrapper tool Trim Galore! to trim reads with low quality bases (Phred score < 30) and Illumina adapter sequences [66]. We used kallisto [67] to pseudoalign reads to the *X. birchmanni* reference transcriptome and imported raw counts for differential gene expression analysis into the R package DESeq2 [68]. Briefly, we created a DESeqDataSet object using the tximport package, setting *X. birchmanni* as the reference group. We performed log-fold change shrinkage using an adaptive shrinkage estimator with a fitted mixture of normal distributions as a prior, derived from the ‘ashr’ package [69]. Counts were normalized to plot expression profiles using DESeq’s internal normalization, which calculates a normalization factor per sample using a median of ratios method. Genes with zero counts, extreme outliers (using a Cook’s distance cutoff of 0.99), or a low mean normalized counts were removed from analysis. To check for potential bias in the pseudoalignment step, we also pseudoaligned trimmed reads against the *X. malinche* reference transcriptome, repeated all analyses, and obtained qualitatively identical results (Supporting Information 5).

### Allele-specific expression analysis

We used a modified version of the program WASP [70] to test for evidence for allele-specific expression of genes differentially expressed between *X. birchmanni* and *X. malinche* in the regenerating caudal tissue of F_1_ hybrids (https://github.com/TheFraserLab/Hornet). WASP corrects for mapping biases that can impact analyses of allele-specific expression by identifying reads that overlap SNPs and removing reads that show evidence of mapping biases. This resulted in counts for the *X. birchmanni* and *X. malinche* alleles at each ancestry informative SNP in F_1_ hybrids. DESeq2 [68] was used to analyze this count data. Counts were imported into a DESeq object as a matrix with the design ∼ individual + allele. All size factors were set to one to avoid size factor normalization, and the model was fit with parametric dispersion.

### Pathway and functional category enrichment analysis in regenerating sword tissue

We investigated evidence for functional enrichment among differentially expressed genes using Gene Ontology and KEGG pathway analysis. For KEGG pathway enrichment, we used *X. birchmanni* vs *X. malinche* regenerating fin log fold changes calculated for DESeq2 differential expression analysis. Gene IDs were mapped to Entrez IDs from the *X. maculatus* Ensembl database (version 99) and KEGG pathway gene sets were generated with the kegg.gsets function from *X. maculatus* KEGG IDs. Both databases were downloaded between 30 March 2020 and 2 April 2020. Enriched gene sets were inferred with the R package gage [71] for both single and dual directionality (Table S8). For GO enrichment, we used BioMart to extract *X. maculatus* Ensembl IDs and generated a gene universe using all genes included in *X. birchmanni* vs *X. malinche* day 10 regeneration DESeq2 analysis. We used a hypergeometric test (hyperGTest in R) to obtain a set of overrepresented GO biological pathway terms in the significantly differentially expressed (FDR adjusted p-value < 0.1) gene set (Table S9).

### Substitution and tolerance predictions at candidate sword genes within the QTL region

For each of the candidates in the associated sword QTL region (Table S3), we generated predicted cDNA alignments based on the *X. birchmanni* and *X. malinche* genomes and available genome sequences for other species [72–74] and quantified rates of amino acid evolution using PAML [75]. Using the known phylogenetic relationships between species [14] we identified derived substitutions in these genes, with a focus on derived substitutions in *X. birchmanni*. We similarly used PAML to test for evidence of differences in evolutionary rates on the branch leading to *X. birchmanni*.

We also compared individual substitutions in *X. birchmanni* and *X. malinche* sequences in detail using SIFT [39]. Using the *X. malinche, X. cortezi, X. montezumae, X. nezahualcoyotl*, and *X. hellerii* (outgroup) sequences, we identified derived amino acid changes in *X. birchmanni*. We then extracted all protein sequences for bony fish from NCBI’s protein database and aligned them with clustal omega [76]. We evaluated this alignment with SIFT and asked whether derived substitutions in *X. birchmanni* were predicted to change protein function.

### Ancestry near the chromosome 13 QTL in natural hybrid populations

To ask whether patterns of *X. malinche* ancestry in natural hybrid populations were unusual in our QTL as a whole and at candidate genes inside the QTL region, we generated joint null distributions for each population. For each hybrid population for which we had previously inferred local ancestry [24,28,43], we generated summaries of average ancestry in 1 Mb and 50 kb windows across the 24 swordtail chromosomes. Next, we generated expected null distributions for ancestry across the four focal populations. We randomly drew a window from each population and recorded the ancestry. For each population, we determined whether the randomly drawn value had equivalent or lower *X. malinche* ancestry than observed in the focal QTL region for that population. We repeated this procedure 5,000 times and asked how frequently randomly drawn ancestry from all four populations was equal to or lower than true *X. malinche* ancestry.

### Phylogenetic approaches

For phylogenetic analyses, we needed sequences from each species of interest aligned to the same coordinate system (Table S7). To generate these sequences, we mapped reads from each species to the *X. birchmanni* reference genome [43] using *bwa* [77]. Next, we removed duplicates with picard tools, realigned indels, called variants using GATK [78], and filtered variants as previously described [24]. We used these variant sites to generate alignments of phylogenetically informative sites for each species on chromosome 13 (https://github.com/Schumerlab/Lab_shared_scripts).

For each gene of interest within the QTL peak, we extracted the alignment, which included both exons and introns, and ran the program RAxML [79] with 100 rapid bootstraps. Following this step, we used RAxML to infer maximum likelihood phylogenies for these regions using the General Reversible Time substitution model. We examined the output for evidence of regions with unusual topologies that received high bootstrap support, which may indicate the presence of incomplete lineage sorting (ILS) or gene flow.

We were also interested in inferring phylogenetic evidence for gene flow between *X. variatus* and *X. birchmanni* and *X. malinche* using this dataset. We used the program PhyloNet-HMM [44] which uses pre-defined hybridization topologies and gene trees to infer local ancestry in the presence of ILS. Past work has shown this approach to have a relatively low false-switching rate in the presence of ILS [74] and our simulations suggest we should have high power to identify introgressed regions (Supporting Information 7). Specifically, we evaluated whether there were regions within the QTL interval on chromosome 13 that supported gene flow from *X. variatus* into *X. birchmanni* but not from *X. variatus* into *X. malinche*, using a posterior probability threshold of 0.9.

## Acknowledgements

We thank Jeremy Berg, Jenn Coughlan, Mark Kirkpatrick, Hakhamanesh Mostafavi, Molly Przeworski, Yuval Simons, Mike Ryan, Andrew Taverner, and members of the Rosenthal and Schumer labs for helpful discussion and/or feedback on earlier versions of this work. We thank the federal government of Mexico for permission to collect fish. Stanford University and the Stanford Research Computing Center provided computational support for this project. This work was supported by NSF LTREB 1354172 to G. G. Rosenthal and a Hanna H. Gray fellowship and NIH 1R35GM133774 grant to M. Schumer.

## Supporting Information Captions

**Fig S1. Sword length by group and QTL effect size estimates. A**. Distribution of normalized sword length phenotypes in F_1_ and F_2_ hybrids between *X. birchmanni* and *X. malinche* and within each of these species. These distributions allow us to estimate broad sense heritability for sword length. **B**. Posterior distribution of ABC simulations to estimate the proportion of phenotypic variance explained by the sword length QTL on chromosome 13. The red line indicates the maximum a posteriori estimate of 0.055. This analysis indicates that the chromosome 13 QTL explains a substantial proportion of the heritable variation in sword length (∼11%) but suggests the presence of other QTL underlying the sword.

**Fig S2. Local ancestry along chromosomes 1-12 for an F**_**2**_ **hybrid individual**. Plotted here are the number of *X. malinche* alleles at each ancestry informative site supported by a posterior probability of 0.9 or greater for a given ancestry state. Scale on x-axis corresponds to the chromosome length in megabases.

**Fig S3. Expected individual level accuracy from simulations of early generation hybrids**. Simulations were conducted using the *mixnmatch* and *ancestryinfer* programs with parameters matching those observed in our study system. See Supporting Information 1 for more details.

**Fig S4. Sword length and sword black margin can become decoupled in hybrids even though the traits are always observed in *X. malinche***. Top - *X. malinche*. Bottom left - male hybrid with a short sword lacking upper and lower pigmented margin. Bottom right - male hybrid with a short sword lacking an upper sword pigmented margin but displaying a lower sword pigmented margin.

**Fig S5. MUMmer alignment of chromosome 13**. This alignment, generated from the *X. birchmanni* and *X. malinche de novo* assemblies, indicates that there are no structural rearrangements between species in the QTL region. The approximate location of the QTL region is indicated by the gray box. Red dots indicate co-linear alignments, blue dots indicate inverted alignments.

**Fig S6. Sword length is associated with genome-wide ancestry**. Sword length is associated with genome-wide ancestry in early generation hybrids between *X. birchmanni* and *X. malinche* (Spearman’s ρ = 0.2, p<4×10^−6^). The correlation between sword length and genome-wide ancestry remains even after accounting for ancestry on chromosome 13 (Spearman’s ρ = 0.18, p<4×10^−5^).

**Fig S7. QTL analysis with and without AIC chromosomes included as covariates**. Thirteen chromosomes were retained in the AIC analysis of the association between chromosome-level ancestry and sword length. We repeated QTL mapping including ancestry on each of these chromosomes as covariates, excluding chromosome 13, and confirmed that we still detect the chromosome 13 QTL in this analysis.

**Fig S8. Stages of sword regeneration**. Example of *X. birchmanni* (left), F_1_ (middle), and *X. malinche* (right) fish included in the sword regeneration RNAseq experiment. Shown here are phenotypes from fish pre-removal of the edge of the caudal fin (and sword in *X. malinche* and F_1_s), post-removal, and after tissue regrowth.

**Fig S9. Heatmap of 30 differentially expressed genes between *X. birchmanni* and *X. malinche* regenerating tissue with the strongest expression differences between species**. 30 out of the 3,333 significantly (FDR adjusted p-value < 0.1) differentially expressed genes are shown. Many of these genes have intermediate expression in F_1_ hybrids. Dark blue, light blue, and red rectangles under the dendrogram indicate species identity of each biological replicate, light blue to orange shading in the matrix indicates relative expression level. Color legend to the right corresponds to log_2_ fold changes in expression.

**Fig S10. Expression of *sp8* in regenerating caudal tissue in *X. malinche* (MM), *X. birchmanni* (BB), and F**_**1**_ **hybrids (MB)**. Semi-transparent points show normalized counts for each individual and solid points and whiskers show the mean ± two standard errors of the mean. The log fold change between *X. birchmanni* and *X. malinche* for *sp8* estimated by DESeq2 was 0.17 (p=0.22).

**Fig S11. Genome wide ancestry distribution in naturally occurring hybrid populations versus average ancestry within the QTL region (dashed line)**. Plotted here is ancestry in 1 Mb windows for the whole genome versus the 1.2 Mb window overlapping with the chromosome 13 QTL peak. Ancestry in this entire region is not unusual compared to the null expectations across hybrid populations, however there is substantial variation in ancestry within the QTL region (see Fig 4).

**Fig S12. Local phylogeny and inferred ancestry at *sp8* with PhyloNet-HMM. A**. Local phylogeny of the *sp8* region generated with RAxML with the GTR+GAMMA model. Node labels show bootstrap support based on 100 rapid bootstraps. **B**. Local ancestry results using PhyloNet-HMM at *sp8* region (dashed gray lines) indicate that although there is evidence of introgression from platyfish nearby, there are no introgressed regions from the platyfish clade that are unique to *X. birchmanni. X. malinche* is shown in red, *X. birchmanni* in blue; purple dots indicate overlap in posterior probability of introgression between the two species.

**Fig S13. QTL results for three chromosomes with (blue) and without (black) genome-wide ancestry included as a covariate**. Without ancestry included as a covariate (black lines), we recover three QTLs that pass the genome-wide significance threshold (LOD=4). However, the peaks on chromosome 1 and chromosome 20 have a relatively flat signal and drop below the genome-wide significance threshold when we account for genome-wide ancestry in R/qtl analysis (blue lines).

**Fig S14. Genome-wide distribution of *X. malinche* ancestry among F**_**2**_ **hybrids in our mapping population**. Although on average individuals derive 50% of their genome from each parental species, there is substantial variance in ancestry generated by the recombination process.

**Fig S15. Results of admix’em simulations evaluating accuracy of heritability estimates varying the level of genetic variation for the trait of interest in parental populations**. Broad sense heritability of a phenotype as a function of ancestry was varied from 0.2 to 0.6 across simulations. We also varied whether all phenotypic variation was attributable to ancestry (blue) or whether there was segregating variation for the phenotype within the simulated *X. malinche* population (red). In those simulations we drew allele frequencies for the causal loci in the *X. malinche* population from a random exponential distribution and arbitrarily set the effect size to 1% of the simulated QTL effect size. Large points and whiskers show mean and two standard deviations across 100 simulations, raw data per simulation is shown by individual points. **B**. In another series of simulations, we varied the effect size of the alleles segregating in the simulated *X. malinche* population from 3-10% of the QTL effect size (red). Simulations with no segregating variation in *X. malinche* are shown in blue for comparison. Large points and whiskers show mean and two standard deviations across 100 simulations, raw data per simulation is shown by individual points. See Supporting Information 10 for a complete description of these simulations.

**Fig S16. Predicted power curve for QTL detection in our study as a function of simulated effect size**. We varied the proportion of phenotypic variation explained by a single QTL and tested our power to detect it in simulations. Each point represents the proportion of 1,000 simulations in which the QTL was detected at our genome-wide significance threshold.

**Fig S17. Results of a two-dimensional two-QTL genome scan for sword length**. Data shown above and left of the diagonal compare a full two QTL model allowing for epistatic interactions to a single QTL model (LOD_fv1_). Results shown below and right of the diagonal show comparison of a two QTL model, where two single loci on separate chromosomes have additive effects on the phenotype, to a single QTL model (LOD_av1_). The color indicates the magnitude of the LOD score. Numbers on the color scale to the right correspond to values of LOD_fv1_ and LOD_av1_, respectively.

**Fig S18. Schematic showing measurement of sword length using ImageJ software**. Sword extension was normalized by standard length for QTL analysis of sword length.

**Fig S19. Manhattan plot showing the results of R/qtl analysis for the presence of the lower sword pigmented edge**. Sword length is one of several phenotypes that makes up the composite sword trait (Fig. S4). The upper sword edge and lower sword edge are also important components of the trait. No genome-wide significant QTL were identified for the lower sword edge trait.

**Fig S20. Expression levels of differentially expressed *hoxa* genes in regenerating caudal fin tissue in *X. malinche* (dark blue), F**_**1**_ **hybrids (light blue) and *X. birchmanni* (red)**. This cluster of genes is found nearbly the QTL region we identify on chromosome 13. Solid points and whiskers show mean and two standard errors of the mean, semi-transparent points show the raw data per individual.

**Fig S21. Example of results from simulations of ancient admixture and application of PhyloNet-HMM to infer local ancestry**. Black dots show the posterior probability inferred by PhyloNet-HMM that the site is hybridization derived. Purple lines above the black dots show the true locations of admixture derived tracts, determined from decoding tree sequences in SLiM.

**Fig S22. Comparison of linear model and R/qtl results**. Linear model based mapping results (right) for sword length mirror results from R/qtl (left). Due to extremely long runtimes it was impractical to use R/qtl in simulations but these results suggest that a linear model based approach gives qualitatively similar results.

**Fig S23. Distribution of median p-values in simulations of polygenic traits determined by ancestry at 50 underlying loci. A**. Distribution of median p-values at ancestry informative sites in each of 100 simulations when genome-wide ancestry is not included as a covariate in the analysis. **B**. Distribution of median p-values at ancestry informative sites in each of 100 simulations when genome-wide ancestry is included as a covariate in the analysis. Red dashed line indicates the median of each distribution. P-values are skewed towards lower values in **A**. This may reflect lower power when genome-wide ancestry is included as a covariate, or reflect the impacts of uncorrected ancestry structure on p-value distributions.

**Fig S24. Example results for simulations of 50 loci contributing to variation in a polygenic trait. A**. Manhattan plot showing results without genome-wide ancestry included as a covariate. Manhattan plot showing results for the same simulation with genome-wide ancestry included as a covariate. Because we should not have power to detect individual QTL in these simulations, the results in **A** reflect possible inflation of associations when genome-wide ancestry is not accounted for. Red line shows genome-wide significance threshold used in our study.

**Fig S25. Distribution of median p-values in simulations of polygenic traits determined by ancestry at 500 underlying loci. A**. Distribution of median p-values at ancestry informative sites in each of 100 simulations when genome-wide ancestry is not included as a covariate in the analysis. **B**. Distribution of median p-values at ancestry informative sites in each of 100 simulations when genome-wide ancestry is included as a covariate in the analysis. Red dashed line indicates the median of each distribution. P-values are significantly shifted towards lower values in **A**. Given that we should have near zero power to detect loci of these effect sizes in our simulations, this skew likely reflects the impacts of uncorrected ancestry structure on p-value distributions.

**Fig S26. Example results for simulations of 500 loci contributing to variation in a polygenic trait. A**. Manhattan plot showing results without genome-wide ancestry included as a covariate. **B**. Manhattan plot for the same simulation showing results with genome-wide ancestry included as a covariate. Because we should not have power to detect individual QTL in these simulations, the results in **A** reflect possible inflation of associations when genome-wide ancestry is not accounted for. Red line shows genome-wide significance threshold used in our study.

**Fig S27. Distribution of average crossover number across 24 chromosomes in the lab-generated hybrids used in this study**. The mean number of crossovers per chromosome in this dataset was 1.3, similar to observed values in F_2_ hybrids (1.2), suggesting that the majority of individuals included in our mapping population were F_2_ hybrids.

**Fig S28. Manhattan plot of R/qtl mapping results for sword length excluding F**_**3**_ **or later individuals**. Individuals that are likely F_3_ hybrids given the number of observed crossover events (Fig S27) were excluded from this analysis. Red line indicates genome-wide significant threshold determined by permutation.

**Table S1. Genes differentially expressed in regenerated sword tissue**. Information on 3,333 significantly differentially expressed genes between *X. birchmanni* and *X. malinche* in the regenerating caudal fin.

**Table S2. All genes that overlap with the joint QTL region**. Genes that fall within the joint QTL interval identified on chromosome 13.

**Table S3. Summary of expression, annotation, and substitution evidence for candidate genes in the QTL interval**. Evidence associated with the strongest candidates within the QTL region on chromosome 13, those associated with fin or limb phenotypes, growth, skeletal or muscle phenotypes.

**Table S4. Summary of evolutionary analyses for top candidates**. SIFT, joint ancestry, and dN/dS analysis at other candidate genes identified in the chromosome 13 QTL region. Joint ancestry analysis was conducted using four hybrid populations and comparing observed *X. malinche* ancestry across populations to that expected from randomly sampling windows for each population.

**Table S5. Summary of ancestry change analyses for top candidates**. Ancestry changes at other candidate genes identified in the chromosome 13 QTL interval over time in Acuapa time series data.

**Table S6. Read information for RNAseq analysis**. Number of reads collected per individual included in RNAseq-based analysis of sword regeneration.

**Table S7. SRA accessions for previously published datasets used in phylogenetic analysis**. Average per basepair coverage when mapped to the *X. birchmanni* reference genome is listed.

**Table S8. KEGG pathway enrichment in significantly differentially expressed genes between *X. malinche* vs *X. birchmanni* regenerating fin tissue**. Only two pathways were enriched for higher expression in *X. birchmanni*, suggesting that either *X. birchmanni* upregulates or *X. malinche* downregulates (or has constitutively lower expression) of genes involved in ECM-receptor interaction and focal adhesion. This analysis included all genes at FDR adjusted p-value < 0.1.

**Table S9. Overrepresented Gene Ontology terms in the significantly differentially expressed genes between *X. malinche* vs *X. birchmanni* regenerating fin tissue**. Of 3368 GO terms tested, 216 terms were found to be significantly overrepresented with a p-value < 0.05. This analysis included all genes at FDR adjusted p-value < 0.1.

**Supporting Information text 1**. Expected accuracy of local ancestry inference

**Supporting Information text 2**. Narrowing the QTL interval

**Supporting Information text 3**. Effect size of the chromosome 13 QTL and expected power

**Supporting Information text 4**. Trait independence in hybrids

**Supporting Information text 5**. Gene expression analysis of regenerating sword tissue **Supporting Information text 6**. Inference of selection on *sp8* region in time transect data **Supporting Information text 7**. Power to detect introgression with PhyloNet-HMM

**Supporting Information text 8**. Simulations of polygenic traits and QTL analysis

**Supporting Information text 9**. Evaluating possible complexity introduced by the cross design

**Supporting Information text 10**. Investigating impacts of genetic variation within *X. malinche* on heritability estimates

## References

1. Kirkpatrick M, Hall DW. Sexual selection and sex linkage. Evolution. 2004;58(4):683–91.

2. Mead LS, Arnold SJ. Quantitative genetic models of sexual selection. Trends Ecol Evol (Amst). 2004 May;19(5):264–71.

3. Morgan MD, Pairo-Castineira E, Rawlik K, Canela-Xandri O, Rees J, Sims D, et al. Genome-wide study of hair colour in UK Biobank explains most of the SNP heritability. Nat Commun. 2018 Dec 10;9(1):1–10.

4. Wood AR, Esko T, Yang J, Vedantam S, Pers TH, Gustafsson S, et al. Defining the role of common variation in the genomic and biological architecture of adult human height. Nat Genet. 2014 Nov;46(11):1173–86.

5. Kunte K, Zhang W, Tenger-Trolander A, Palmer DH, Martin A, Reed RD, et al. doublesex is a mimicry supergene. Nature. 2014 Mar;507(7491):229–32.

6. Jones FC, Grabherr MG, Chan YF, Russell P, Mauceli E, Johnson J, et al. The genomic basis of adaptive evolution in threespine sticklebacks. Nature. 2012 Apr;484(7392):55–61.

7. Linnen CR, Poh Y-P, Peterson BK, Barrett RDH, Larson JG, Jensen JD, et al. Adaptive evolution of multiple traits through multiple mutations at a single gene. Science. 2013 Mar 15;339(6125):1312–6.

8. Lamichhaney S, Fan G, Widemo F, Gunnarsson U, Thalmann DS, Hoeppner MP, et al. Structural genomic changes underlie alternative reproductive strategies in the ruff (Philomachus pugnax). Nat Genet. 2016 Jan;48(1):84–8.

9. Lampert KP, Schmidt C, Fischer P, Volff J-N, Hoffmann C, Muck J, et al. Determination of onset of sexual maturation and mating behavior by melanocortin receptor 4 polymorphisms. Curr Biol. 2010 Oct 12;20(19):1729–34.

10. Kim K-W, Jackson BC, Zhang H, Toews DPL, Taylor SA, Greig EI, et al. Genetics and evidence for balancing selection of a sex-linked colour polymorphism in a songbird. Nat Comm. 2019 Apr 23;10(1):1852.

11. Rockman MV. The Qtn program and the alleles that matter for evolution: all that’s gold does not glitter. Evolution. 2012;66(1):1–17.

12. Xu S. Theoretical basis of the Beavis Effect. Genetics. 2003 Dec 1;165(4):2259–68.

13. Jones JC, Fan S, Franchini P, Schartl M, Meyer A. The evolutionary history of Xiphophorus fish and their sexually selected sword: a genome-wide approach using restriction site-associated DNA sequencing. Mol Ecol. 2013;22(11):2986–3001.

14. Cui R, Schumer M, Kruesi K, Walter R, Andolfatto P, Rosenthal G. Phylogenomics reveals extensive reticulate evolution in Xiphophorus fishes. Evolution. 2013;67(8):2166–2179.

15. Basolo AL. A further examination of preexisting bias favoring a sword in the genus Xiphophorus. Anim Behav. 1995;50:365–75.

16. Basolo AL. Female preference predates the evolution of the sword in swordtail fish. Science. 1990;250(4982):808–10.

17. Rauchenberger M, Kallman KD, Morizot DC. Monophyly and geography of the Rio Panuco Basin Mexico swordtails genus Xiphophorus with descriptions of four new species. Am Mus Novit. 1990;(2975).

18. Basolo AL. Female preference for male sword length in the green swordtail, Xiphophorus helleri (Pisces: Poeciliidae). Anim Behav. 1990;40(2):332–8.

19. Delclos PJ, Forero SA, Rosenthal GG. Divergent neurogenomic responses shape social learning of both personality and mate preference. J Exp Biol. 2020;223(6).

20. Cui R, Delclos PJ, Schumer M, Rosenthal GG. Early social learning triggers neurogenomic expression changes in a swordtail fish. Proc Biol Sci. 2017 May 17;284(1854).

21. Verzijden MN, Culumber ZW, Rosenthal GG. Opposite effects of learning cause asymmetric mate preferences in hybridizing species. Behav Ecol. 2012 Sep 1;23(5):1133–9.

22. Lande R. Models of speciation by sexual selection on polygenic traits. PNAS. 1981 Jun 1;78(6):3721–5.

23. Wong BBM, Rosenthal GG. Female disdain for swords in a swordtail fish. Am Nat. 2006;167(1).

24. Schumer M, Xu C, Powell DL, Durvasula A, Skov L, Holland C, et al. Natural selection interacts with recombination to shape the evolution of hybrid genomes. Science. 2018 May 11;360(6389):656.

25. Culumber ZW, Fisher HS, Tobler M, Mateos M, Barber PH, Sorenson MD, et al. Replicated hybrid zones of Xiphophorus swordtails along an elevational gradient. Mol Ecol. 2011;20(2):342–56.

26. Rosenthal GG, de la Rosa Reyna XF, Kazianis S, Stephens MJ, Morizot DC, Ryan MJ, et al. Dissolution of sexual signal complexes in a hybrid zone between the swordtails Xiphophorus birchmanni and Xiphophorus malinche (Poeciliidae). Copeia. 2003;2003(2):299–307.

27. Falconer DS, Mackay TFC. Introduction to quantitative genetics. 4th ed. Harlow: Addison Wesley Longman; 1996.

28. Schumer M, Powell DL, Delclós PJ, Squire M, Cui R, Andolfatto P, et al. Assortative mating and persistent reproductive isolation in hybrids. Proc Natl Acad Sci USA. 2017 Oct 10;114(41):10936.

29. Basolo A. Shift in investment between sexually selected traits: tarnishing of the silver spoon. Anim Behav. 1998 Mar;55(3):665–71.

30. Schumer M, Powell DL, Corbett-Detig R. Versatile simulations of admixture and accurate local ancestry inference with mixnmatch and ancestryinfer. bioRxiv. 2019 Nov 30;860924.

31. Broman KW, Wu H, Sen Ś, Churchill GA. R/qtl: QTL mapping in experimental crosses. Bioinformatics. 2003 May 1;19(7):889–90.

32. Schartl M, Kneitz S, Ormanns J, Schmidt C, Anderson JL, Amores A, et al. The developmental and genetic architecture of the sexually selected male ornament of swordtails. bioRxiv. 2020.

33. Otto SP, Jones CD. Detecting the undetected: estimating the total number of loci underlying a quantitative trait. Genetics. 2000 Dec 1;156(4):2093–107.

34. Basolo AL, Trainor BC. The conformation of a female preference for a composite male trait in green swordtails. Anim Behav. 2002 Mar 1;63(3):469–74.

35. Offen N, Blum N, Meyer A, Begemann G. Fgfr1 signalling in the development of a sexually selected trait in vertebrates, the sword of swordtail fish. BMC Devel Biol. 2008;8.

36. Offen N, Meyer A, Begemann G. Identification of novel genes involved in the development of the sword and gonopodium in swordtail fish. Devel Dyn. 2009;238(7):1674–87.

37. Shibata E, Yokota Y, Horita N, Kudo A, Abe G, Kawakami K, et al. Fgf signalling controls diverse aspects of fin regeneration. Development. 2016 Aug 15;143(16):2920–9.

38. Kawakami Y, Esteban CR, Matsui T, Rodríguez-León J, Kato S, Belmonte JCI. Sp8 and Sp9, two closely related buttonhead-like transcription factors, regulate Fgf8 expression and limb outgrowth in vertebrate embryos. Development. 2004 Oct 1;131(19):4763–74.

39. Vaser R, Adusumalli S, Leng SN, Sikic M, Ng PC. SIFT missense predictions for genomes. Nat Protoc. 2016 Jan;11(1):1–9.

40. Völler D, Reinders J, Meister G, Bosserhoff A-K. Strong reduction of AGO2 expression in melanoma and cellular consequences. Br J Cancer. 2013 Dec 10;109(12):3116–24.

41. Li Y-H, Chen H-Y, Li Y-W, Wu S-Y, Wangta-Liu, Lin G-H, et al. Progranulin regulates zebrafish muscle growth and regeneration through maintaining the pool of myogenic progenitor cells. Sci Rep. 2013 Jan 31;3(1):1–8.

42. Tayebi N, Jamsheer A, Flöttmann R, Sowinska-Seidler A, Doelken SC, Oehl-Jaschkowitz B, et al. Deletions of exons with regulatory activity at the DYNC1I1 locus are associated with split-hand/split-foot malformation: array CGH screening of 134 unrelated families. Orphanet J Rare Dis. 2014 Jul 29;9:108.

43. Powell DL, García-Olazábal M, Keegan M, Reilly P, Du K, Díaz-Loyo AP, et al. Natural hybridization reveals incompatible alleles that cause melanoma in swordtail fish. Science. 2020 May 15;368(6492):731–6.

44. Liu KJ, Dai J, Truong K, Song Y, Kohn MH, Nakhleh L. An HMM-based comparative genomic framework for detecting introgression in eukaryotes. PLOS Comput Biol. 2014 Jun 12;10(6):e1003649.

45. Patterson N, Moorjani P, Luo Y, Mallick S, Rohland N, Zhan Y, et al. Ancient admixture in human history. Genetics. 2012 Nov 1;192(3):1065–93.

46. Darwin C. the descent of man, and selection in relation to sex. D. Appleton; 1871. 508 p.

47. Treichel D, Schöck F, Jäckle H, Gruss P, Mansouri A. mBtd is required to maintain signaling during murine limb development. Genes Dev. 2003 Nov 1;17(21):2630–5.

48. Bell SM, Schreiner CM, Waclaw RR, Campbell K, Potter SS, Scott WJ. Sp8 is crucial for limb outgrowth and neuropore closure. PNAS. 2003 Oct 14;100(21):12195–200.

49. Draper BW, Stock DW, Kimmel CB. Zebrafish fgf24 functions with fgf8 to promote posterior mesodermal development. Development. 2003 Oct;130(19):4639–54.

50. Orr HA. The population genetics of adaptation: the distribution of factors fixed during adaptive evolution. Evolution. 1998;52(4):935–49.

51. Fisher RA. The genetical theory of natural selection. иипол Классик; 289 p.

52. Simon A, Bierne N, Welch JJ. Coadapted genomes and selection on hybrids: Fisher’s geometric model explains a variety of empirical patterns. Evol Lett. 2018;2(5):472–98.

53. Rieseberg LH, Archer MA, Wayne RK. Transgressive segregation, adaptation and speciation. Heredity. 1999 Oct;83(4):363–72.

54. Visscher PM, Haley CS. Detection of putative quantitative trait loci in line crosses under infinitesimal genetic models. Theor Appl Genet. 1996 Oct;93(5–6):691–702.

55. Pfaff CL, Parra EJ, Bonilla C, Hiester K, McKeigue PM, Kamboh MI, et al. Population structure in admixed populations: effect of admixture dynamics on the pattern of linkage disequilibrium. Am J Hum Genet. 2001 Jan;68(1):198–207.

56. Wiens JJ. Widespread loss of sexually selected traits: how the peacock lost its spots. Trends Ecol Evol. 2001 Sep 1;16(9):517–23.

57. Rosenthal GG, Flores Martinez TY, García de León FJ, Ryan MJ. Shared preferences by predators and females for male ornaments in swordtails. Am Nat. 2001 Aug;158(2):146–54.

58. Achorn AM, Rosenthal GG. It’s not about him: mismeasuring ‘good genes’ in sexual selection. Trends Ecol Evol. 2020 Mar 1;35(3):206–19.

59. Endler JA. Signals, signal conditions, and the direction of evolution. The Am Nat. 1992 Mar 1;139:S125–53.

60. Rosenthal G. Mate choice: the evolution of sexual decision making from microbes to humans. Princeton University Press; 648 p.

61. Kallman KD, Bao IY. Female heterogamety in the swordtail, Xiphophorus alvarezi Rosen (Pisces, Poeciliidae), with comments on a natural polymorphism affecting sword coloration. J Exp Zool. 1987 Jul;243(1):93–102.

62. Schneider CA, Rasband WS, Eliceiri KW. NIH image to imagej: 25 years of image analysis. Nat Methods. 2012 Jul;9(7):671–5.

63. Corbett-Detig R, Nielsen R. A Hidden Markov Model approach for simultaneously estimating local ancestry and admixture time using next generation sequence data in samples of arbitrary ploidy. PLOS Genet. 2017 Jan 3;13(1):e1006529.

64. Falconer DS. Introduction to quantitative genetics. Oliver & Boyd, Edinburgh & London.; 1960. ix + 365 pp.

65. Marçais G, Delcher AL, Phillippy AM, Coston R, Salzberg SL, Zimin A. MUMmer4: A fast and versatile genome alignment system. PLOS Comput Biol. 2018 Jan 26;14(1):e1005944.

66. Krueger F. FelixKrueger/TrimGalore. Available from: https://github.com/FelixKrueger/TrimGalore

67. Bray NL, Pimentel H, Melsted P, Pachter L. Near-optimal probabilistic RNA-seq quantification. Nat Biotechnol. 2016 May;34(5):525–7.

68. Love MI, Huber W, Anders S. Moderated estimation of fold change and dispersion for RNA-seq data with DESeq2. Genome Biol. 2014 Dec 5;15(12):550.

69. Stephens M. False discovery rates: a new deal. Biostatistics. 2017 Apr 1;18(2):275–94.

70. van de Geijn B, McVicker G, Gilad Y, Pritchard JK. WASP: allele-specific software for robust molecular quantitative trait locus discovery. Nat Methods. 2015 Nov;12(11):1061–3.

71. Luo W, Friedman MS, Shedden K, Hankenson KD, Woolf PJ. GAGE: generally applicable gene set enrichment for pathway analysis. BMC Bioinformatics. 2009;10(1):161.

72. Schartl M, Walter RB, Shen Y, Garcia T, Catchen J, Amores A, et al. The genome of the platyfish, Xiphophorus maculatus, provides insights into evolutionary adaptation and several complex traits. Nat Genet. 2013 Mar 31;45:567.

73. Shen Y, Chalopin D, Garcia T, Boswell M, Boswell W, Shiryev SA, et al. X. couchianus and X. hellerii genome models provide genomic variation insight among Xiphophorus species. BMC Genomics. 2016 Jan 7;17(1):37.

74. Schumer M, Cui R, Powell DL, Rosenthal GG, Andolfatto P. Ancient hybridization and genomic stabilization in a swordtail fish. Mol Ecol. 2016 Jun 1;25(11):2661–79.

75. Yang Z. PAML 4: phylogenetic analysis by maximum likelihood. Mol Biol Evol. 2007 Aug 1;24(8):1586–91.

76. Sievers F, Wilm A, Dineen D, Gibson TJ, Karplus K, Li W, et al. Fast, scalable generation of high-quality protein multiple sequence alignments using Clustal Omega. Mol Syst Biol. 2011 Oct 11;7:539.

77. Li H, Durbin R. Fast and accurate short read alignment with Burrows-Wheeler transform. Bioinformatics. 2009;25(14).

78. McKenna A, Hanna M, Banks E, Sivachenko A, Cibulskis K, Kernytsky A, et al. The Genome Analysis Toolkit: a MapReduce framework for analyzing next-generation DNA sequencing data. Genome Res. 2010 Sep;20(9):1297–303.

79. Stamatakis A. RAxML-VI-HPC: Maximum likelihood-based phylogenetic analyses with thousands of taxa and mixed models. Bioinformatics. 2006;22(21):2688–90.

