## Supporting Information for "The genetic architecture of the sexually selected sword ornament and its evolution in hybrid populations"

### Supporting Information 1. Expected accuracy of local ancestry inference

We performed simulations using the *mixnmatch* pipeline [1] to evaluate the expected accuracy of local ancestry inference for $F_2$ hybrids between *X. birchmanni* and *X. malinche*. *mixnmatch* simulates admixed genomes and outputs a table of ancestry tracts for each individual as well as simulated Illumina reads. For the purposes of these simulations we provided sequence information and the local recombination map from chromosome 1 of *X. birchmanni*. As in the real data we modeled two generations of admixture, 0.2X coverage, and simulated cross-well contamination rates of 2% [1]. We inferred local ancestry using the *ancestryinfer* pipeline [1] which implements the hidden Markov model applied to our real data [2]. We summarized accuracy by comparing the true simulated ancestry to the inferred ancestry state, using a posterior probability threshold of 0.9.

Results of these simulations indicate that we expect local ancestry calls for $F_2$ hybrids to be highly accurate (Fig. S3), with estimated per-site error rates of <0.2%. This expectation corresponds well to observed ancestry patterns in $F_2$ hybrids, where overall admixture proportions and the number of ancestry transitions per chromosome mirror our expectations for early generation hybrids (Fig. S2).

### Supporting Information 2. Narrowing the QTL interval

#### *Supporting Information 2.1. Generating a confidence interval for the QTL region*

Because this project relied on a mapping population of early generation hybrids, the 1.5 LOD interval associated with the sword QTL on chromosome 13 was large, spanning 4.1 Mb and containing 186 genes (13.2 to 17.3 Mb). We evaluated whether there was sufficient information to narrow this region further by bootstrapping our data. Because we found R/qtl to be prohibitively slow for use in simulations, even when analyzing a single chromosome, we first verified that results were nearly identical using a linear-model based approach (see Supporting Information 8) and then proceeded using this approach in simulations.

We subsampled our dataset to 75% the total number of individuals in 1,000 replicate simulations. For each simulation we recorded the location of the peak associated marker on chromosome 13. This allowed us to evaluate the extent to which we expect the peak to move as a result of sampling noise in the data. The 95% confidence intervals of these locations (15.3-16.9 Mb) yielded a 1.7 Mb region at the center of the QTL we had initially identified.

#### *Supporting Information 2.2. Results of an independent mapping study of sword length in X. hellerii and X. maculatus*

While analyzing our results, we discovered that another research group had independently conducted an experiment to map the genetic basis of sword length by crossing sworded (*X. hellerii*) and swordless (*X. maculatus*) species. These two species are deeply diverged from *X. birchmanni* and *X. malinche* (1.6-2% sequence divergence), and from each other (~2% sequence divergence). Using a similar QTL mapping approach, they generated $F_1$ hybrids between *X. hellerii* males and *X. maculatus* females and second-generation backcross hybrids by crossing $F_1$s with *X. hellerii* females [3,4]. Individuals were genotyped with a RADseq approach [3] and the relationship between SNP genotype and sword length was analyzed using R/qtl. They identified a QTL overlapping with the region we identified on chromosome 13, with the 1.5 LOD interval matching the region from 14.3 to 16.5 Mb in the *X. birchmanni* assembly [5]. At first glance, this suggests that the genetic architecture of the sword phenotype is shared in these two species pairs and that comparisons between studies could be used to further narrow the region of interest.

However, since QTL intervals identified by the two studies were relatively large, we wanted to evaluate the probability that both studies would have identified the same region by chance. To do so, we permuted a 2.2 Mb region onto the genome 1,000 times and asked how frequently it overlapped by chance with the QTL interval that we identified by bootstrapping. We found that overlap of the two QTL was unexpected by chance (p-value by permutation 0.004). This suggests that the overlap of the two QTL is not coincidental and that we can use the joint signal in our efforts to identify causal genes within the QTL region. Thus, in subsequent analyses we focus on the narrower region identified from both mapping studies, the interval from 15.3 to 16.5 Mb on chromosome 13.

*Supporting Information 2.3 Identifying candidate genes in the associated interval on chromosome 13*

The 1.2 Mb region identified by the analyses discussed above contained a total of 52 genes. We next performed a series of analyses to identify the genes in this interval most likely to be responsible for the QTL association.

From this subset of 52 genes, we first asked which had annotations known to be associated with limb or tail phenotypes, growth phenotypes, or skeletal phenotypes based on aggregated data in the GeneCards database [6], narrowing our focal gene list to sixteen (Table S3). Next, we asked which genes were expressed in regenerating sword tissue in *X. malinche* and $F_1$ hybrids. We reasoned that genes that are not expressed during sword regeneration are unlikely to be involved in the sword length trait, leaving us with fourteen focal genes (Table S3).

Of this subset of 14 genes, we predicted that to drive the QTL signal, they should be responsible for an effect in *cis*. This could be accomplished either by an amino acid difference between *X. birchmanni* and *X. malinche* that underlies the morphological difference between species, differences in expression between the two species, or some combination of the two. We thus evaluated which of these genes had nonsynonymous amino acid changes between species and which had expression differences in regenerating caudal fin tissue between *X. malinche* and *X. birchmanni*. This analysis led to a set of 8 possible candidates which are listed along with their annotation information and sequence and expression results in Table S3.

**Supporting Information 3. Effect size of the chromosome 13 QTL and expected power**

*Supporting Information 3.1. Simulations to infer the effect size of the chromosome 13 QTL*

We wanted to know how much of the variation in sword length is explained by the significant QTL on chromosome 13. Based on the coefficient in a linear model, we estimated that the chromosome 13 QTL explains approximately 5% of the total variation in sword length, or ~10% of the heritable variation in length. However, we knew that it was possible that this effect size was an overestimate due to a well-known statistical phenomenon where effect sizes for QTL detected in studies with low power are often inflated (known as the Beavis effect or the winner's curse [7]).

The inflated effect size estimates associated with the winner's curse are the result of enforcing genome-wide significance thresholds in scenarios in which there is low power to detect the true signal at that threshold. Regions of the genome where noise in the data matches the direction of the underlying signal are more likely to pass the significance threshold. We thus sought to infer the likely effect size of the chromosome 13 QTL, using an approximate Bayesian approach. Importantly, by implementing the same genome-wide significance threshold as used with our real data in simulations we should capture the impacts of effect size inflation due to the winner's curse, allowing us to more accurately estimate effect size of the chromosome 13 QTL region.

We extracted the observed genotypes for each individual at the QTL peak, generated simulated phenotypes, and performed mapping analysis. We used the following steps to do so in simulations:

1) We drew the proportion of phenotypic variance explained by the chromosome 13 QTL ($chr13_{effect}$) from a uniform prior that ranged from zero to our empirical heritability estimate for sword length (0.48).
    a. Using these values and the average sword phenotype in our mapping population, we partitioned the phenotypic variance contributed by chromosome 13 ($pheno_{chr13}$) and the variance contributed by the rest of the genome and by environmental factors ($pheno_{other}$).
2) We next generated simulated sword phenotypes for each individual.
    a. Based on an individual's genotype at the QTL peak on chromosome 13, we performed the following steps:
        i. If an individual was homozygous *X. malinche*, we assigned that individual a phenotype value of $pheno_{chr13}$.
        ii. If an individual was heterozygous, we assigned that individual a phenotype of half of $pheno_{chr13}$.
        iii. If an individual was homozygous *X. birchmanni* we assigned that individual zero for $pheno_{chr13}$.
    b. We next added trait variation to mimic variation explained by the rest of the genome and environmental sources.
        i. We drew from a normal distribution with mean equal to $pheno_{other}$, and the variance set to the observed variance in sword length in $F_2$ hybrids.
        ii. We added $pheno_{chr13}$ to $pheno_{other}$ to generate a simulated sword length for each individual.
3) We performed a linear regression analysis as we had done for the real data and determined the p-value for the association between chromosome 13 and simulated sword length, as well as the proportion of the phenotypic variance explained by the chromosome 13 QTL. We performed rejection sampling based on the observed p-value in the real data, using a tolerance of 5%.
4) This procedure was repeated until 5,000 simulations had been selected.

These simulations resulted in a well-resolved posterior distribution of possible effect sizes for the chromosome 13 QTL (Fig. S1). Surprisingly, we found the maximum a posteriori estimate of effect size was not lower than our initial estimate of effect size for this QTL (5.5% compared to point estimate of 5%; 95 % CIs: 0.05-0.12). This suggests that the QTL on chromosome 13 falls within the range of effect sizes that we have power to detect in our study.

*Supporting Information 3.2. Simulations to infer the power threshold for QTL mapping*

In addition to inferring the effect size of the chromosome 13 QTL, we were interested in determining the threshold at which we have power to detect QTL in our study. We see ample evidence that there are other loci contributing to variation in sword length and inferring the likely effect size at which we have power to detect QTL gives us some information about the QTL we were unable to map.

We performed simulations following a similar procedure to those described above except that instead of using an ABC framework, we performed a grid search to ask how frequently simulated QTL of a given effect size were detected. The effect sizes in the grid ranged from 0-50% of the phenotypic variance explained in increments of 5%. For each value, we performed simulations using the following steps:

1) Using observed genome-wide ancestry for each individual in our mapping population, we generated simulated diploid genotypes.
    a. We drew from a random binomial distribution with probability equal to the proportion of *X. malinche* ancestry genome-wide in that individual.
2) Using the effect size drawn for the simulation and the average sword phenotype in our mapping population, we partitioned the phenotypic variance contributed by the simulated QTL ($pheno_{QTL}$) and that contributed by the rest of the genome and environmental factors ($pheno_{other}$).
3) We next generated simulated sword phenotypes for each individual.
    a. Based on an individual's genotype at the simulated QTL peak, we performed the following steps:
        i. If an individual was homozygous *X. malinche*, we assigned that individual a phenotype value of $pheno_{QTL}$.
        ii. If an individual was heterozygous, we assigned that individual a phenotype of half of $pheno_{QTL}$.
        iii. If an individual was homozygous *X. birchmanni* we assigned that individual zero for $pheno_{QTL}$.
    b. We next added trait variation to mimic variation explained by the rest of the genome and environmental sources.
        i. We drew from a normal distribution with mean equal to $pheno_{other}$, and the variance set to the observed variance in sword length in $F_2$ hybrids.
        ii. We added $pheno_{QTL}$ to $pheno_{other}$ to generate a simulated sword length for each individual.
4) We performed QTL analysis at the simulated QTL as we had in the real data, including genome-wide ancestry as a covariate, and asked if the p-value exceeded the genome-wide significance threshold used in our mapping analyses.

5) For each effect size, we repeated this procedure 1,000 times and recorded the proportion of simulations in which the simulated QTL was detected.

The results of these simulations, which are shown in Fig. S16, allow us to approximate the effect size at which we have power to detect underlying QTL given our sample size. They suggest that we have power to detect QTL that explain ~5% of the heritable variation in about 50% of simulations (i.e. $\theta$ in Otto & Jones [8]) and QTL that explain ~10% of the heritable variation in 90% of simulations. QTL that explain a smaller degree of trait variation (~1%) are rarely detected. Thus, we expect our design is missing many QTL that contribute to variation in sword length. We note that the results of these simulations are consistent with those described in *Supporting Information 3.1* since they predict that we have approximately 90% power to detect a QTL of the effect size we infer.

*Supporting Information 3.3. Exploring the possible number of causal loci for sword length*

Although the model fitting results we present in the main text suggest a lower bound for the number of genes underlying the sword phenotype, we wanted to explore the likely number of loci contributing to phenotypic variation in the sword further. We used existing theory to infer the possible number of QTL underlying sword length [8]. Specifically, we used the approach of Otto & Jones [8] to calculate the likely number of QTL underlying a trait, given the observed heritable difference in the focal trait between the parental species ($D$), the average effect size of identified QTL ($M$), and the effect size at which we have power to detect QTL in our study ($\theta$). This approach assumes an exponential distribution of QTL effect sizes and additive effects of the QTL [8]. We note that because we only detected one genome-wide significant QTL, these estimates should be viewed with caution, as they are strongly dependedent on the inferred effect size of that QTL.

We have an estimate of $M$ for the QTL on chromosome 13 from a linear regression and the ABC simulations described above. We also indirectly estimated $\theta$ in power simulations, by determining the effect size at which we have at least 50% power to detect a QTL given our study design. Importantly, the estimated $M$ value for the chromosome 13 QTL is only slightly higher than estimated $\theta$, again highlighting the likelihood of many missed QTL for sword length.

The results of this calculation suggest that we could expect as many as ~150 loci underlying sword length (Fig. 2; Materials & Methods, Supporting Information 3). However, we emphasize that uncertainty in effect sizes, assumptions about the underlying distribution of effect sizes, and estimates that rely on a single detected QTL limit our confidence in this analysis.

*Supporting Information 3.4. Results of multi-QTL model analysis*

Acknowledging that our initial mapping likely missed many regions of the genomes contributing to variation in sword length, we performed a two-dimensional QTL scan for sword length, with hybrid index as a covariate using the scantwo function in R/qtl. We used the same 12,794 markers as we previously used in the one-dimensional scan (see main text; [9]) but calculated conditional QTL genotype probabilities using the calc.geneoprob() function on a coarser grid (step =3). As in the one-dimensional scan we performed interval mapping using the EM algorithm. Recombination fraction was estimated using the est.rf function and markers missing genotype data were excluded using the drop.nullmarker function. The threshold for genome-wide likelihood of odds (LOD) at false discovery rates of 5% and 10% was determined based on 1,000 permutations of sword phenotypes onto the observed genotypes. We did not identify a two-locus model with a significantly better fit to the data than a simpler one-locus model even at the more permissive 10% false discovery rate (Fig. S17).

### Supporting Information 4. Trait independence in hybrids

We phenotyped artificial hybrids and parental individuals for three different traits related to the sword phenotype: sword length, the upper pigmented sword margin, and the lower pigmented sword margin (Fig. S18; Materials & Methods). We used a partial correlation approach to test the extent to which these traits were correlated with each other in our mapping panel.

We found that the sword upper edge phenotype was strongly correlated with sword length ($R=0.48$, $p<10^{-32}$), and thus chose not to map this as a separate trait. By contrast, sword lower edge was not as strongly predictive of sword length ($R= 0.19$, $p<10^{-5}$). We thus performed QTL mapping for the sword lower edge as described in the main text (Materials & Methods) except that we modeled a binary trait. Surprisingly, we did not find evidence for any genome-wide significant QTL associated with the sword lower margin (Fig. S19), nor a clear association with genome-wide ancestry ($R=0.05$, $p=0.26$).

### Supporting Information 5. Gene expression analysis of regenerating sword tissue

#### *Supporting Information 5.1. Gene expression analysis using the X. malinche reference transcriptome*

For our primary expression analysis, we use the *X. birchmanni* transcriptome. However, we were concerned about the potential for downstream biases in expression results generated by relying on a reference sequence from a single species. To control for this, we quantified count abundance with kallisto [10] using both the *X. birchmanni* and *X. malinche* reference transcriptomes, and performed all downstream differential expression analysis steps in parallel.

As expected, log fold change values were strongly correlated across analyses using the two different reference sequences, with an adjusted $R^2$ of 0.92 (based on a linear model in R). Moreover, of 3,333 and 3,374 significantly differentially expressed genes identified using the *X. birchmanni* and *X. malinche* references, respectively, 3190 genes were identified in both analyses. This suggests minimal impacts of choice of reference sequence on our results. Results of downstream analyses were also similar. GO enrichment yielded 227 significantly overrepresented terms for the analysis using the *X. birchmanni* reference, and 214 terms for the analysis using the *X. malinche* reference, with 202 terms identified in both analyses. KEGG enrichment results were identical between analyses.

#### *Supporting Information 5.2. Expression signal of hox genes in regenerating sword tissue*

The QTL we identify on chromosome 13 occurs nearby a cluster of *hoxa* genes. Because of their well-known role in regulating the animal body plan, we were initially intrigued by this result. However, the *hoxa* cluster does not overlap with the narrower interval we identify using bootstrapping, nor does it overlap with the QTL region identified in crosses between *X. maculatus* and *X. hellerii*. Nevertheless, given the known role of *hox* genes, we investigated expression patterns at individual *hoxa* genes.

We found that four of the seven genes in the *hoxa* cluster were differentially expressed in regenerating caudal fin tissue between *X. birchmanni* and *X. malinche* at an adjusted p-value threshold of 0.05. $F_1$s tended to have intermediate expression levels (Fig. S20). Because many genes are differentially expressed between *X. birchmanni* and *X. malinche* in regenerating caudal tissue, we asked how unusual it was for four out of a randomly drawn set of seven genes to show significant expression differences. We generated a null distribution by sampling seven random genes and recorded whether four or more of those genes were differentially expressed at an adjusted p-value of 0.05. We found that the pattern observed at the *hoxa* cluster is unexpected by chance (p=0.005 based on 1,000 permutations). However, we caution that due to confounding factors such as correlations in expression between neighboring genes [11] and non-independence of *hoxa* genes due to shared regulatory networks [12], this analysis may not be conservative.

*Supporting Information 5.3. Concordance between differential and allele-specific expression during sword regeneration in $F_1$ hybrids*

For $F_1$ hybrids, we can also ask which of the expression changes we observe between the parental species are in part attributable to evolved changes in *cis*. Although we only identified two genes with significant allele specific expression out of the 3,926 that passed our quality thresholds, we evaluated whether there was evidence for similar patterns in the allele specific and differential expression datasets. For all genes that were differentially expressed between *X. birchmanni* and *X. malinche* in regenerating sword tissue and present in our allele specific expression data, we evaluated whether there was concordance between allelic expression ratios in $F_1$s and overall direction of expression level differences between species.

We extracted genes that were differentially expressed between *X. birchmanni* and *X. malinche* at a corrected p-value threshold of 0.05 and their log fold changes. For these genes, we extracted estimated fold changes in expression of the *X. birchmanni* and *X. malinche* alleles in $F_1$s, generated from WASP and DEseq2 ([13], 724 genes present in both datasets). We asked, using a paired sign test, whether the allelic expression ratios were enriched for the observed direction of expression differences between species. We found evidence of substantial enrichment ($S=503$, $p<10^{-16}$). In addition, we found a moderate correlation in the fold change estimated for differential expression and allele specific expression using this approach ($\rho= 0.24$, $p=10^{-10}$).

*Supporting Information 5.4. Lack of correlation between chromosome-level expression patterns and estimated effect size*

Given that ancestry on more than a dozen chromosomes predicts sword phenotype in hybrids and that thousands of genes are differentially expressed between *X. birchmanni* and *X. malinche* in regenerating fin tissue (see Fig. 2, Fig. 3), we asked whether there was a relationship between them. Specifically, we asked whether chromosomes with larger estimated effects on sword length had a greater number (or proportion) of differentially expressed genes. We did not detect a relationship between the estimated chromosome-level effect sizes and number (or proportion) of differentially expressed genes, regardless of whether we performed the analysis based on all differentially expressed genes, or those with *X. malinche* or *X. birchmanni* biased expression (all comparisons $p > 0.26$).

**Supporting Information 6. Inference of selection on *sp8* region in time transect data**

We find lower than expected *X. malinche* ancestry at the *sp8* gene in aggregate across hybrid populations (Fig. 4). Moreover, in one population where we have access to time series data (the Acuapa population), we observe a significant decrease in *X. malinche* ancestry over time at the *sp8* gene (Fig. 4; two proportions z-test of 2018 versus 2006 – $p = 0.007$).

This decrease in *X. malinche* ancestry over time could be driven by selection or by genetic drift. To explore whether the decline in *X. malinche* ancestry at *sp8* in this time series data was consistent with selection against *X. malinche* ancestry we used an approximate Bayesian approach. We grounded our simulations in observed data from the Acuapa population.

For each simulation, we drew parameters and performed simulations using the following procedure:

1. First, we determined starting parameters for each simulation
    a. We drew a starting ancestry frequency, $f_m$, for 2006 from a uniform distribution. We defined this uniform distribution as ranging from 0.19 to 0.37 *X. malinche* ancestry. This range represents the observed *X. malinche* ancestry at *sp8* in our 2006 sample from Acuapa, plus or minus the standard error of that estimate.
    b. We drew a selection coefficient, $s$, from a uniform distribution ranging from 0.5 to -0.5.
    c. We drew dominance of the *X. malinche* allele, $h$, from a uniform distribution ranging from zero to one.
    d. The number of generations of selection, $g$, was drawn from a uniform distribution ranging from 12 to 36 generations. This is equivalent to the lower and upper bound of the plausible number of generations that could have elapsed during the 12 year sampling period [14].
    e. We drew the diploid hybrid population size, $n$, from a uniform distribution ranging from 200 to 10,000.
2. For each set of parameters, we iterated through $g$ generations of selection
    a. We calculated the predicted *X. malinche* allele frequency in the next generation with selection using the general selection model.
    b. We simulated genetic drift by sampling $2*n$ alleles from a binomial distribution with the probability set to the *X. malinche* frequency in the next generation predicted by the general selection model.
3. We accepted simulations where the final ancestry fell within one standard error of the average ancestry at *sp8* observed in 2018. We repeated simulations until 1,000 parameter sets were accepted.

These simulations resulted in a well-resolved posterior distribution of the strength of selection against *X. malinche* ancestry at *sp8* with a maximum a posteriori (MAP) estimate of -0.1 (95% confidence intervals: -0.44 – -0.03). Estimates of the dominance coefficient, $h$, were skewed towards zero (MAP – 0.01) but 95% confidence intervals were broad (0.007 – 0.94). Posterior distributions for the remaining parameters, including population size, starting ancestry proportions, and number of generations elapsed mirrored the prior distribution. Overall, these simulation results are consistent with moderate selection against *X. malinche* ancestry at *sp8* in the Acuapa population.

Another possible cause for reduced *X. malinche* ancestry not captured in the simulations described above is migration from a population with greater *X. birchmanni* ancestry. Several lines of evidence argue against this. First, we do not observe a significant shift in genome-wide ancestry from 2006-2018 (p=0.1, average *malinche* ancestry 2006=0.26, average *malinche* ancestry 2018=0.29). Second, of 346 individuals sampled from Acuapa over this time period, we did not detect a single ancestry outlier for *X. birchmanni*-like ancestry, but sampled three outliers for *X. malinche*-like ancestry, suggesting that if migration is occurring, it would drive in the opposite direction of our signal.

**Supporting Information 7. Power to detect introgression with PhyloNet-HMM**

*X. birchmanni* lost the sword ornament since it diverged from its common ancestor with *X. malinche*. We wanted to ask whether ancient introgression from another swordless species could explain this, or whether the loss occurred through another mechanism.

*X. variatus* is the only unsworded species that is currently sympatric with *X. birchmanni*, and we previously detected evidence of gene flow from a lineage related to *X. variatus* into *X. birchmanni* and *X. malinche* [15]. We note that we cannot exclude the possibility of different historical distributions of unsworded species.

Because of the deep evolutionary divergence between *X. birchmanni* and the platyfish clade (Fig. 1; containing *X. variatus*), we predicted that we might have power to identify introgression along the genome, even for ancient hybridization events. Specifically, contemporary populations of *X. birchmanni* and *X. variatus* have an average of 1.6% pairwise sequence divergence, or 16 ancestry informative differences per kilobase.

To evaluate our power to detect local introgression from a swordless platyfish species into *X. birchmanni*, we simulated divergence and admixture using SLiM [16], decoded ancestry and genotypes at ancestry informative sites using the tree sequence recording functions in SLiM and pySLiM [17]. We applied a phylogenetic hidden Markov model to genotypes derived from SLiM to identify potentially introgressed tracts [18] and compared these results to the true ancestry.

For SLiM simulations, we used a burn-in of 90,000 generations and simulated four populations with split times equivalent to estimated split times between *X. hellerii*, *X. variatus*, *X. birchmanni*, and *X. malinche*, assuming a historical effective population size of 25,000 [15]. Based on estimates from an $F_4$ ratio test implemented in Admixtools (see below; [19]) we simulated a contribution of 4% of the genome from the lineage leading to *X. variatus* and 96% from the lineage leading to *X. birchmanni*. Although we have found evidence of possible ongoing gene flow between *X. variatus* and *X. birchmanni*, we assumed that a pulse of admixture occurred immediately after the split between *X. birchmanni* and *X. malinche*. This assumption is conservative because ancient gene flow will result in small ancestry tracts that are more difficult to detect using HMM-based methods. Moreover, some of these tracts will also have accumulated mutations since the time of admixture. Thus, this corresponds to the scenario in which we expect to have the lowest power to detect introgressed regions.

We simulated 5 Mb sequences using the recombination map from the first 5 Mb of *X. birchmanni* chromosome 2. We sampled a single individual from each population, converted vcf files generated by SLiM to phylip files using the tool vcf2phylip (https://github.com/edgardomortiz/vcf2phylip), filtered ambiguous sites, and ran the PhyloNet-HMM program [18]. We selected sites that had greater than 0.9 posterior probability support for a given ancestry state and determined whether PhyloNet-HMM inferred the correct ancestry state in that region. We also asked about the false negative rate (i.e. the rate with which tracts are missed). This procedure was repeated 50 times.

The results of these simulations suggest that even in this scenario of ancient admixture we expect to have relatively good power to detect introgressed tracts, likely due to the deep divergence between *X. birchmanni* and species in the platyfish clade. Across simulations, the accuracy per ancestry informative site was 96% and we detected 97.5% of introgressed tracts (Fig. S21).

As a secondary approach, we also calculated the $F_4$ ratio statistic [19] in 1,000 SNP windows. Because the $F_4$ ratio statistic simply relies on site counts it may be more sensitive to short ancestry tracts that are too small to be detected by HMM-based approaches. We used the configuration *X. maculatus X. hellerii* : *X. birchmanni X. malinche* :: *X. maculatus X. hellerii X. variatus X. malinche* implemented through the admixtools program [19]. We again did not find evidence for higher *X. variatus* contribution to the *X. birchmanni* genome near the chromosome 13 QTL peak.

### Supporting Information 8. Simulations of polygenic traits and QTL analysis

#### *Supporting Information 8.1. Impact of including genome-wide ancestry as a covariate in mapping studies*

In R/qtl analysis, we performed mapping with and without genome-wide ancestry included as a covariate. Because we found that excluding genome-wide ancestry as a covariate resulted in two additional QTLs passing our genome-wide significance threshold, we were interested in evaluating the impact of model choice further. Since R/qtl is prohibitively slow for use in even a moderate number of simulations, we instead compared results of linear models conducted in R, after first confirming that in the empirical data these approaches produced nearly identical results (Fig. S22).

To explore the expected impact of including or excluding genome-wide ancestry in the case of moderately to highly polygenic traits, we performed simulations. For each simulation, we used the observed genotypes from our 536 early generation hybrids and simulated phenotypes based on ancestry at 50 underlying loci.

To generate simulated phenotypes, we used the following procedure:

1) First, we partitioned the expected trait variance in hybrids into the environmental and genetic components. We randomly selected 50 sites and treated these as causal loci underlying variation in sword length.
    a. For each individual, to specify the environmental variation in the trait, we drew from a random normal distribution with a mean equal to the trait mean in $F_1$s and variance set to the trait variance observed in $F_1$s.
2) Next, we divided the heritable variation equally among all 50 loci contributing to genetic variation in sword length $F_2$s and generated phenotypes based on simulated genotypes at those loci.

a. If an individual was homozygous for *X. malinche* ancestry at that locus, we added the full effect of that locus to the individual's quantitative phenotype.
b. If an individual was heterozygous at that locus, we added half of the effect of that locus to the individual's quantitative phenotype.
c. If an individual was homozygous *X. birchmanni* at that locus we did not change the phenotype.
d. We repeated this for all 50 loci, generating a polygenic trait for mapping in simulations.

3) We scanned for associations between genotype at each ancestry informative site and simulated phenotype using a linear model, with or without genome-wide ancestry included as a covariate.
4) We performed 100 replicates of these simulations.

Even in simulations with only 50 causal loci, we expect to have very low power to detect individual QTL with our sample of 536 individuals (Fig. S16). We compared p-value distributions for simulated markers in a model that included genome-wide ancestry as a covariate and one that did not. We found that in analyses excluding genome-wide ancestry as a covariate, median (KS test p-value $10^{-9}$; Fig. S23) and minimum p-values were consistently lower (KS test p-value $p<0.001$), contributing to the appearance of peaks of association in some simulations (Fig. S24). This suggests that mapping experiments that do not include genome-wide ancestry as a covariate may have a higher false positive rate when traits are highly polygenic, as has been reported previously [20].

One possible explanation for this observation is that even though power is predicted to be low with our sample size and 50 causal loci, incorporating genome-wide ancestry will further reduce power, and as a result make us less likely to detect true causal loci. We thus repeated the simulations as described above but split the genetic effects over 500 loci and replicated these results (KS test p-value $10^{-7}$; Fig. S25). In the uncorrected simulations we continue to see qualitative results similar to the broader peaks observed in the uncorrected analysis of the real data (Fig. S26). Together, these results suggest that incorporating genome-wide ancestry in simulations will reduce false positives when analyzing traits with a polygenic basis.

*Supporting Information 8.2. Correlations between X. birchmanni ancestry and sword length*

We also used the simulations described above as an opportunity to investigate whether the negative correlations we observe between sword length and *X. malinche* ancestry on a chromosomal level are expected by chance. Specifically, we used the simulations described above and applied the same model selection approach we used in analyzing the real data. For each simulation, we asked whether *X. birchmanni* ancestry was retained in the final model as a predictor of simulated sword length.

While in reality, our simulations only used *X. malinche* ancestry to generate the simulated sword phenotype, we found that in 40% of simulations *X. birchmanni* ancestry was retained as predictive in the final model on at least one chromosome. However, fewer chromosomes contributed to this pattern than in the real data ($p<0.01$ by simulation), and effect sizes estimated for *X. birchmanni* ancestry were uniformly stronger in the real data than any observed in simulations ($p<0.01$ by simulation).

These results could suggest the presence of transgressive effects of *X. birchmanni* ancestry on sword length, or be explained by variations in genetic architecture that we have not yet explored. For example, particular linkage scenarios or certain distributions of QTL effect size could generate stronger correlations between *X. birchmanni* ancestry and sword length, without *X. birchmanni* ancestry itself driving these correlations.

**Supporting Information 9. Evaluating possible complexity introduced by the cross design**

Due to the difficulty of raising sufficient numbers of hybrids in lab, we used a design where we seeded large mesocosm tanks with $F_1$ hybrids (see Materials & Methods). The majority of adults collected within two years after seeding were likely to be $F_2$s, given generation and maturation times. However, a possible drawback of this design is that given some variation in generation and maturation time, sampled individuals could be a mixture of $F_2$ and $F_3$ individuals.

We decided to evaluate this based on comparisons of the number of observed crossovers in sampled individuals to the number expected in $F_2$ hybrids. Specifically, from previous work we had access to local ancestry data from 139 $F_2$ hybrids generated in the lab [1]. We compared the distribution of ancestry transitions, reflecting observed recombination events, between known $F_2$ hybrids and the individuals from our mapping populations (Fig. S27). The median number of crossovers per chromosome was slightly higher in our mapping population (1.3 crossovers/chromosome) than expected from known $F_2$s (1.2 crossovers/chromosome), but the distribution suggests that few $F_3$ or later individuals were included in our mapping population (Fig. S27).

Out of an abundance of caution, we re-ran R/qtl excluding five individuals with numbers of crossovers outside of the range of the distribution of crossovers observed in known $F_2$s. Reassuringly, excluding these individuals had no effect on our results (Fig. S28).

**Supporting Information 10. Investigating impacts of genetic variation within *X. malinche* on heritability estimates**

The approach that we use to estimate heritability is a quantitative genetics based method originally designed for estimating heritability in inbred lines where variation in the parental lines is due solely to environmental variation. While this is likely true in *X. birchmanni* where variance in the caudal fin extension is much less than observed in $F_1$ hybrids, it may not be a valid assumption for *X. malinche*, where there is more variation in sword length among individuals (Fig. S1). Based on comparisons of the phenotypes of *X. birchmanni*, *X. malinche*, and hybrids (Fig. S1), we can infer that the majority of variance in sword length in hybrids is due to ancestry. However, we wanted to explore what impact genetic variation for sword length in *X. malinche* would have on our estimates of ancestry-based heritability.

To do so, we used a simulation approach. We did not attempt to mimic exact parameters from our system, as many of these values are unknown, but rather asked whether adding additional genetic variation in one of the parental populations generally biased estimates of trait heritability as a function of ancestry.

We used the admixture simulator admix'em [21] to simulate genotypes and phenotypes for two parental populations, $F_1$ and $F_2$ hybrids. We simulated 10 chromosomes each with a randomly placed QTL contributing to trait variation. Each QTL explained 10% of the heritable variation in the trait generated by ancestry from the simulated *X. malinche* parental population.

We also varied the amount of environmental variation by adding values from a random uniform distribution such that broad sense heritability due to ancestry was 0.2, 0.4, and 0.6 in three sets of simulations. For each heritability value, we performed two types of simulations. In the first, all heritability was attributable to *X. malinche* ancestry. In the second, segregating polymorphisms at the QTL (implemented in admix'em as loci 1 bp away) could also increase simulated sword length. Allele frequencies for these loci within the simulated *X. malinche* population were drawn from a random exponential distribution. We arbitrarily assigned these loci 1% of the effect size of the QTL they were linked to, but relax this assumption below.

For each value of broad sense heritability, we performed 100 replicate simulations with and without additional phenotypic variation attributable to segregating loci within the simulated *X. malinche* population. Based on these simulations, additional phenotypic variation from *X. malinche* did not appear to impact the accuracy of estimates of ancestry heritability using methods intended for inbred lines. Simulations with and without segregating variation in *X. malinche* resulted in overlapping estimates for heritability attributable to ancestry variation (Fig. S15). Moreover, this observation held when we increased the effect size of the loci segregating in *X. malinche* from 1% of the QTL effect size to 5 and 10% (Fig. S15). We hypothesize that this observation is due to the fact that loci segregating within *X. malinche* contribute to phenotypic variance in both $F_1$ and $F_2$ hybrids, and thus this variance tends to be absorbed in the environmental effect term, resulting in an accurate estimate for heritability attributable to ancestry variation.

**Supplementary Figures**

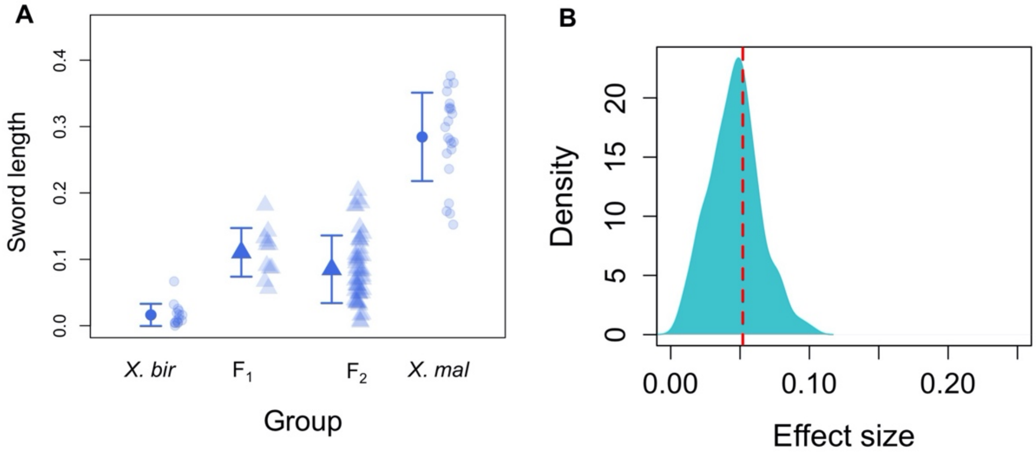


**Fig S1. Sword length by group and QTL effect size estimates. A**. Distribution of normalized sword length phenotypes in $F_1$ and $F_2$ hybrids between *X. birchmanni* and *X. malinche* and within each of these species. These distributions allow us to estimate broad sense heritability for sword length. **B**. Posterior distribution of ABC simulations to estimate the proportion of phenotypic variance explained by the sword length QTL on chromosome 13. The red line indicates the maximum a posteriori estimate of 0.055. This analysis indicates that the chromosome 13 QTL explains a substantial proportion of the heritable variation in sword length (~11%) but suggests the presence of other QTL underlying the sword.

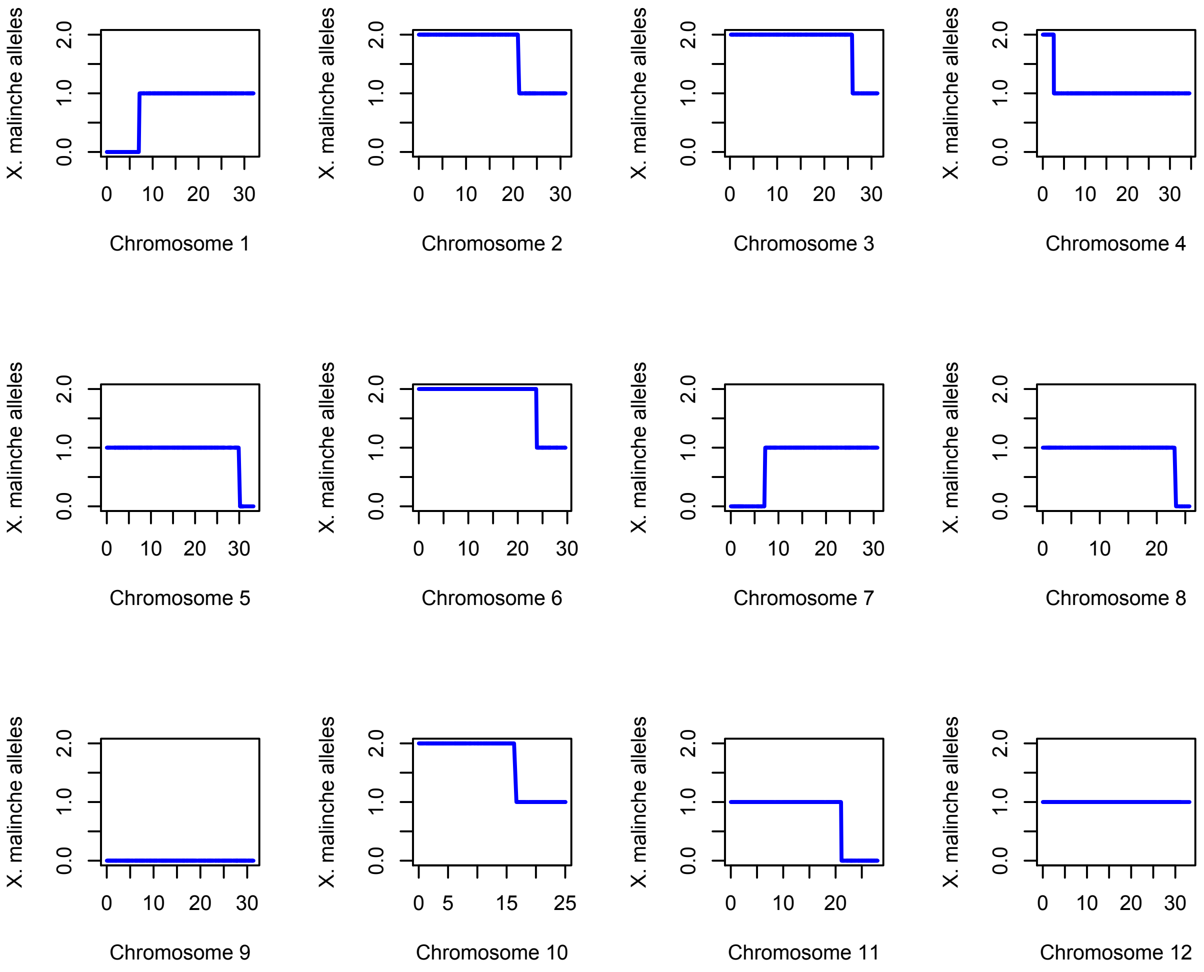


**Fig S2. Local ancestry along chromosomes 1-12 for an $F_2$ hybrid individual.**
Plotted here are the number of *X. malinche* alleles at each ancestry informative site supported by a posterior probability of 0.9 or greater for a given ancestry state. Scale on x-axis corresponds to the chromosome length in megabases.

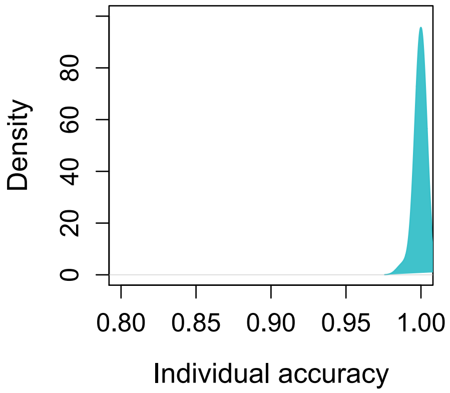


**Fig S3**. **Expected individual level accuracy from simulations of early generation hybrids.** Simulations were conducted using the *mixnmatch* and *ancestryinfer* programs with parameters matching those observed in our study system. See Supporting Information 1 for more details.

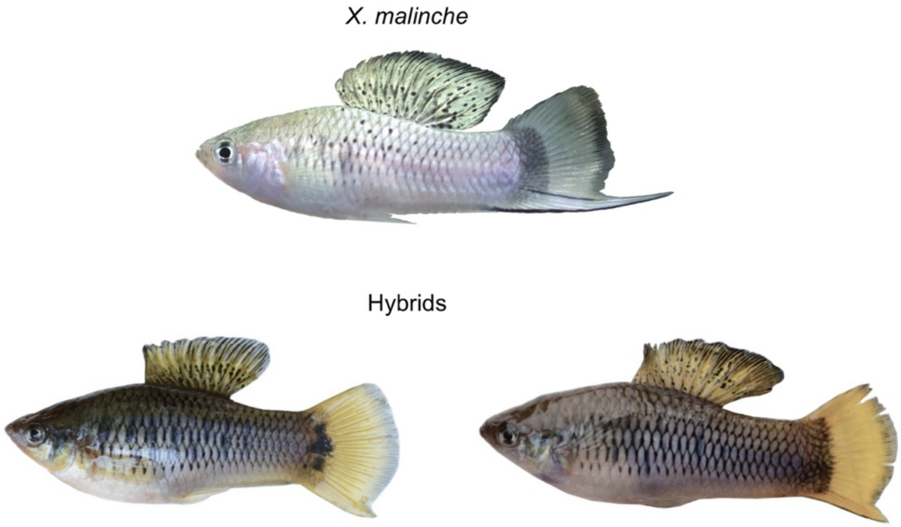


**Fig S4. Sword length and sword black margin can become decoupled in hybrids even though the traits are always observed in *X. malinche*.** Top - *X. malinche.* Bottom left - male hybrid with a short sword lacking upper and lower pigmented margin. Bottom right - male hybrid with a short sword lacking an upper sword pigmented margin but displaying a lower sword pigmented margin.

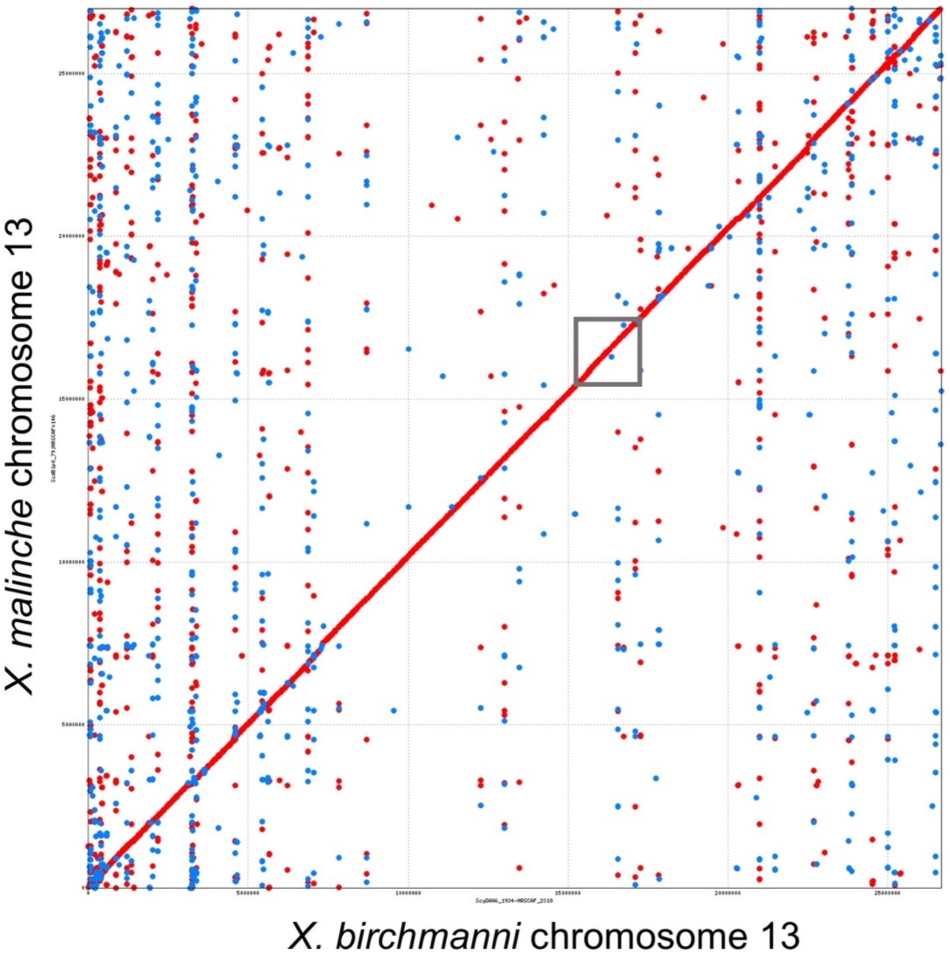


**Fig S5. MUMmer alignment of chromosome 13.** This alignment, generated from the *X. birchmanni* and *X. malinche de novo* assemblies, indicates that there are no structural rearrangements between species in the QTL region. The approximate location of the QTL region is indicated by the gray box. Red dots indicate co-linear alignments, blue dots indicate inverted alignments.

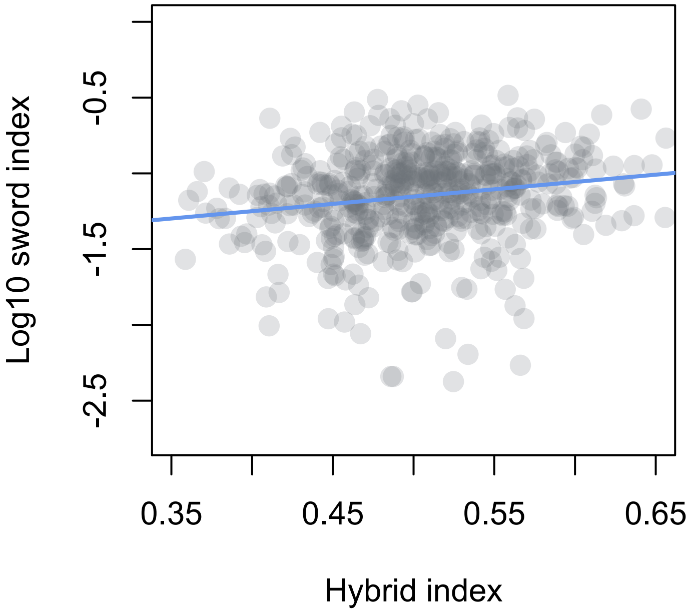


**Fig S6. Sword length is associated with genome-wide ancestry**. Sword length is associated with genome-wide ancestry in early generation hybrids between *X. birchmanni* and *X. malinche* (Spearman's $\rho = 0.2$, $p<4x10^{-6}$). The correlation between sword length and genome-wide ancestry remains even after accounting for ancestry on chromosome 13 (Spearman's $\rho = 0.18$, $p<4x10^{-5}$).

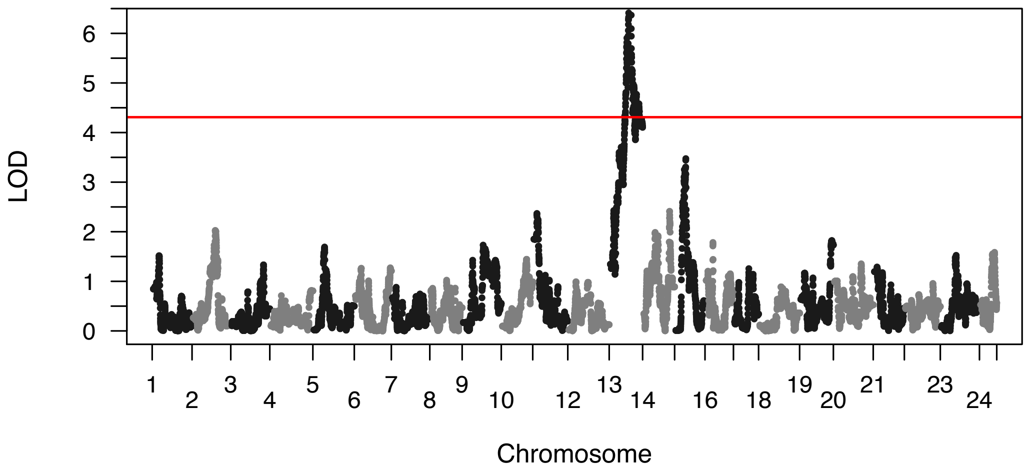


**Fig S7. QTL analysis with and without AIC chromosomes included as covariates.** Thirteen chromosomes were retained in the AIC analysis of the association between chromosome-level ancestry and sword length. We repeated QTL mapping including ancestry on each of these chromosomes as covariates, excluding chromosome 13, and confirmed that we still detect the chromosome 13 QTL in this analysis.

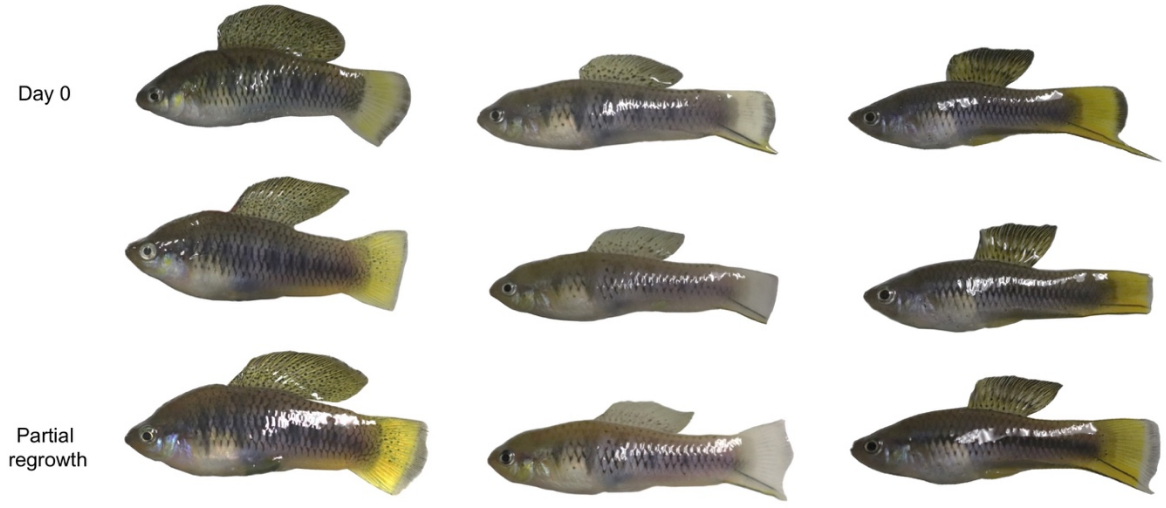


**Fig S8. Stages of sword regeneration.** Example of *X. birchmanni* (left), $F_1$ (middle), and *X. malinche* (right) fish included in the sword regeneration RNAseq experiment. Shown here are phenotypes from fish pre-removal of the edge of the caudal fin (and sword in *X. malinche* and $F_1$s), post-removal, and after tissue regrowth.

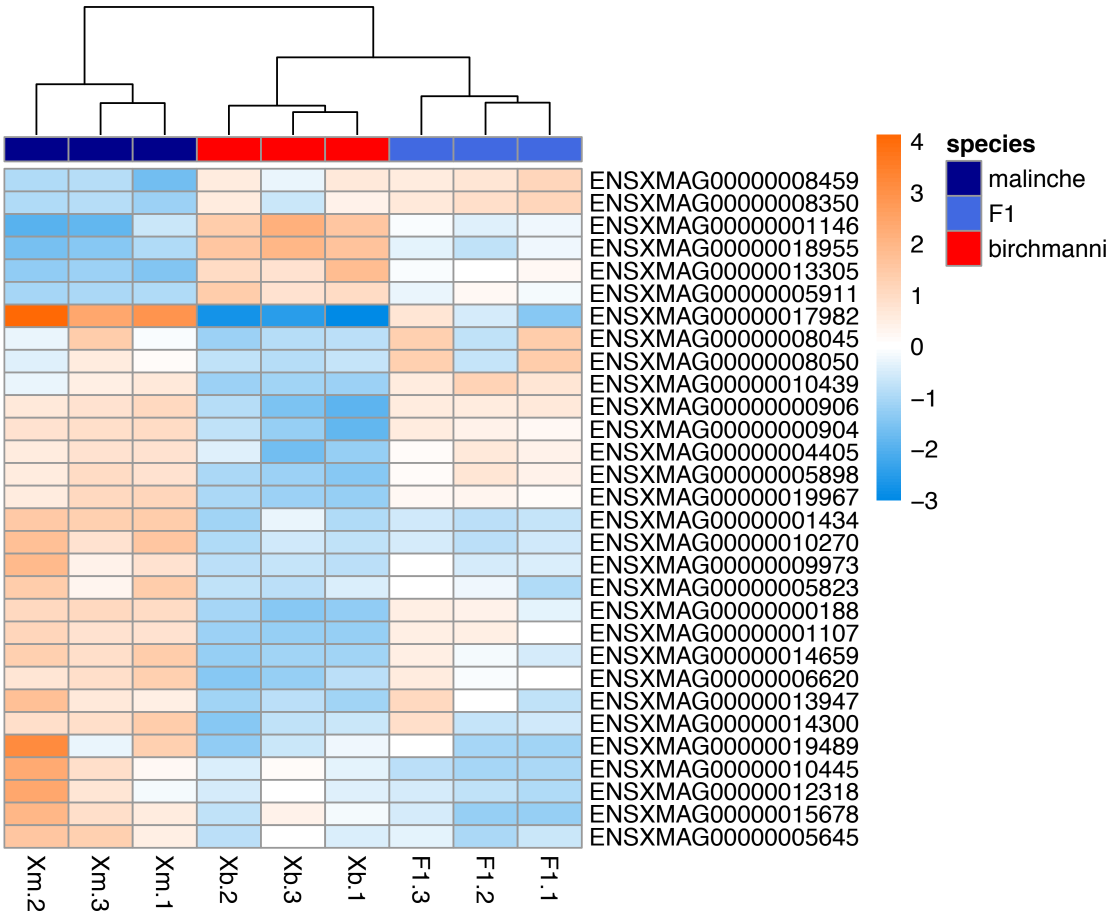


**Fig S9. Heatmap of 30 differentially expressed genes between *X. birchmanni* and *X. malinche* regenerating tissue with the strongest expression differences between species.**
30 out of the 3,333 significantly (FDR adjusted p-value < 0.1) differentially expressed genes are shown. Many of these genes have intermediate expression in $F_1$ hybrids. Dark blue, light blue, and red rectangles under the dendrogram indicate species identity of each biological replicate, light blue to orange shading in the matrix indicates relative expression level. Color legend to the right corresponds to $log_2$ fold changes in expression.

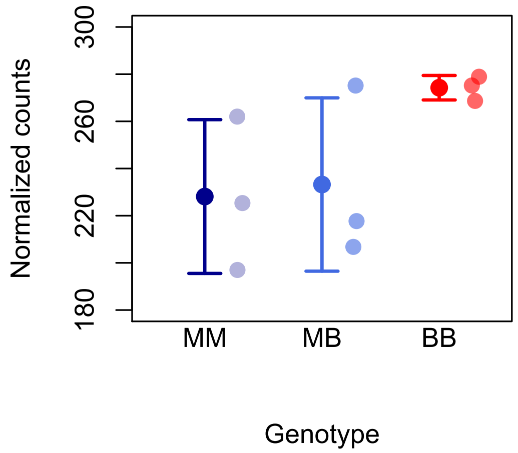


**Fig S10. Expression of *sp8* in regenerating caudal tissue in *X. malinche* (MM), *X. birchmanni* (BB), and $F_1$ hybrids (MB).** Semi-transparent points show normalized counts for each individual and solid points and whiskers show the mean ± two standard errors of the mean. The log fold change between *X. birchmanni* and *X. malinche* for *sp8* estimated by DESeq2 was 0.17 (p=0.22).

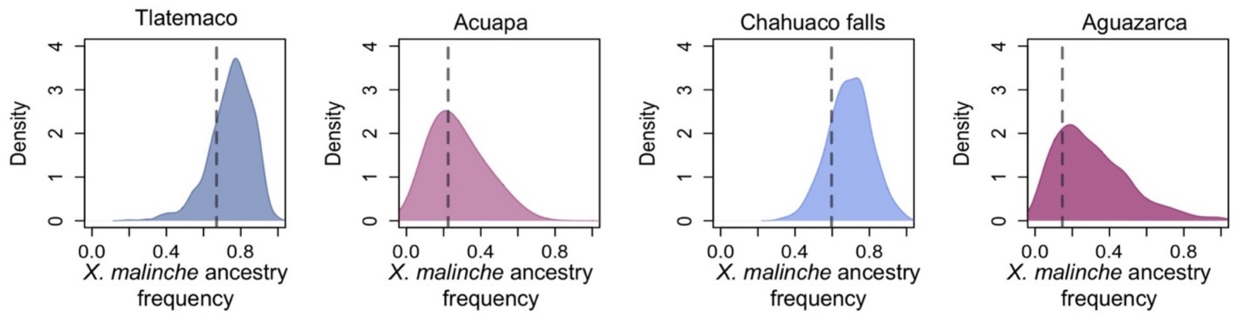


**Fig S11. Genome wide ancestry distribution in naturally occurring hybrid populations versus average ancestry within the QTL region (dashed line).**
Plotted here is ancestry in 1 Mb windows for the whole genome versus the 1.2 Mb window overlapping with the chromosome 13 QTL peak. Ancestry in this entire region is not unusual compared to the null expectations across hybrid populations, however there is substantial variation in ancestry within the QTL region (see Fig 4).

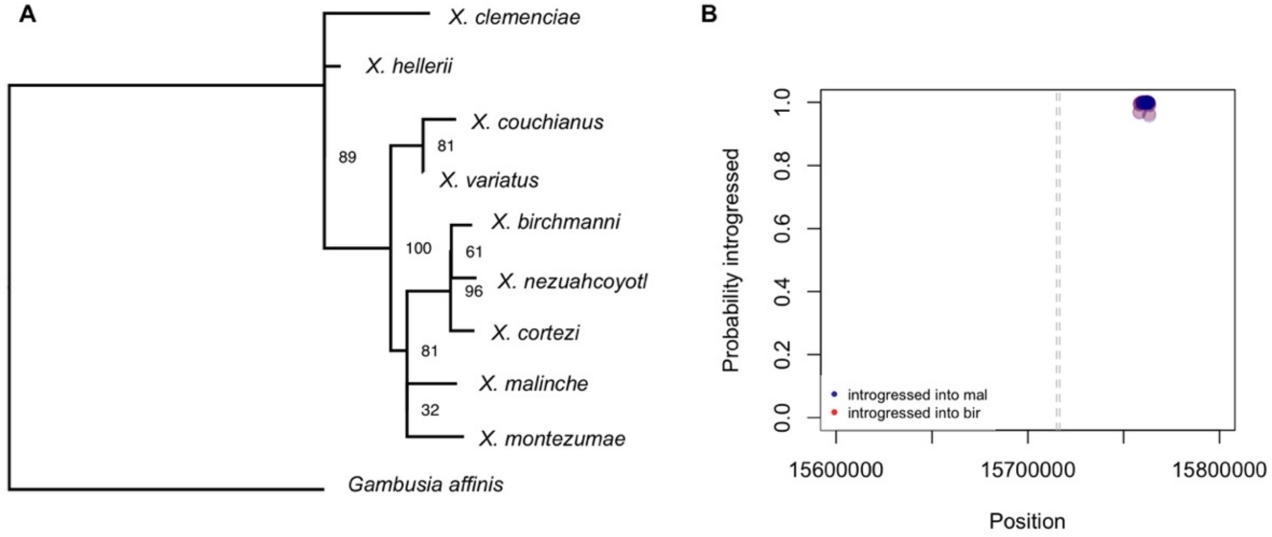


**Fig S12. Local phylogeny and inferred ancestry at *sp8* with PhyloNet-HMM. A.** Local phylogeny of the *sp8* region generated with RAxML with the GTR+GAMMA model. Node labels show bootstrap support based on 100 rapid bootstraps. **B.** Local ancestry results using PhyloNet-HMM at *sp8* region (dashed gray lines) indicate that although there is evidence of introgression from platyfish nearby, there are no introgressed regions from the platyfish clade that are unique to *X. birchmanni*. *X. malinche* is shown in red, *X. birchmanni* in blue; purple dots indicate overlap in posterior probability of introgression between the two species.

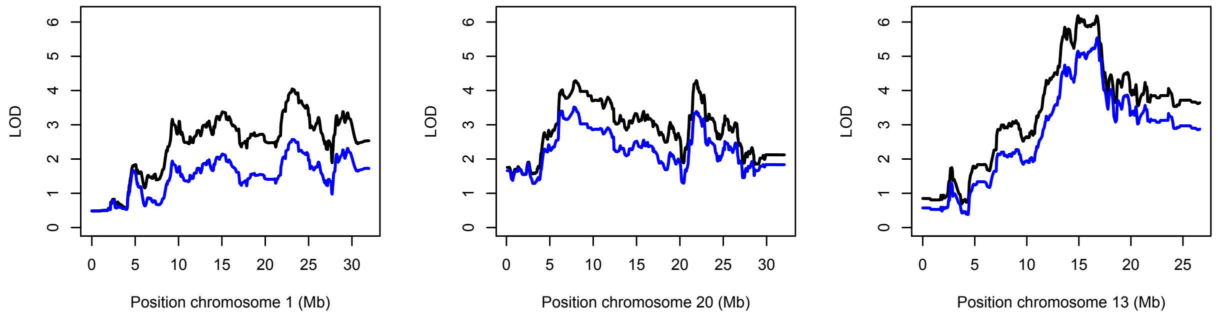


**Fig S13. QTL results for three chromosomes with (blue) and without (black) genome-wide ancestry included as a covariate.** Without ancestry included as a covariate (black lines), we recover three QTLs that pass the genome-wide significance threshold (LOD=4). However, the peaks on chromosome 1 and chromosome 20 have a relatively flat signal and drop below the genome-wide significance threshold when we account for genome-wide ancestry in R/qtl analysis (blue lines).

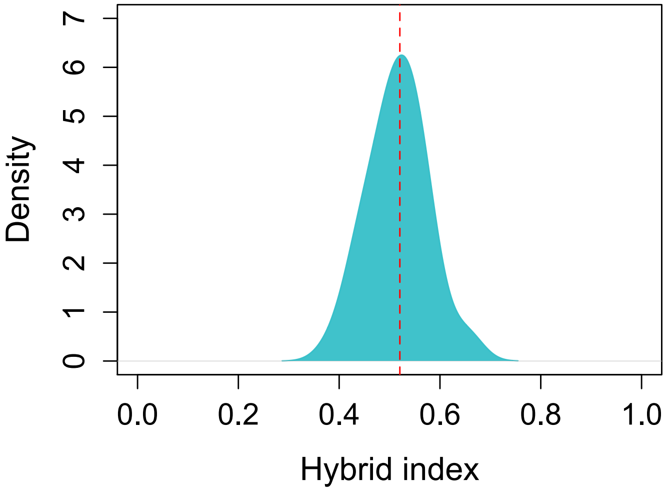


**Fig S14. Genome-wide distribution of *X. malinche* ancestry among $F_2$ hybrids in our mapping population.** Although on average individuals derive 50% of their genome from each parental species, there is substantial variance in ancestry generated by the recombination process.

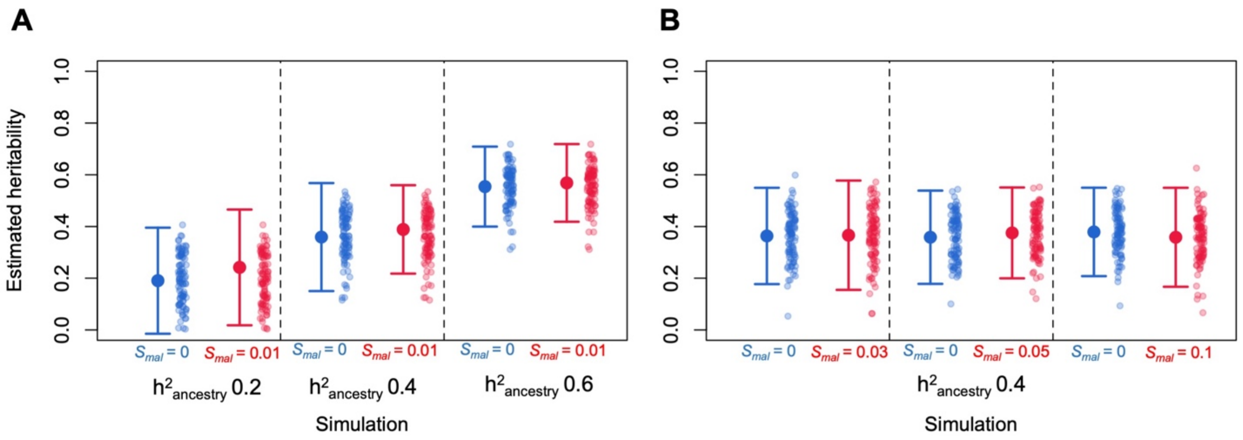


**Fig S15. Results of admix'em simulations evaluating accuracy of heritability estimates varying the level of genetic variation for the trait of interest in parental populations.**
**A**. Broad sense heritability of a phenotype as a function of ancestry was varied from 0.2 to 0.6 across simulations. We also varied whether all phenotypic variation was attributable to ancestry (blue) or whether there was segregating variation for the phenotype within the simulated *X. malinche* population (red). In those simulations we drew allele frequencies for the causal loci in the *X. malinche* population from a random exponential distribution and arbitrarily set the effect size to 1% of the simulated QTL effect size. Large points and whiskers show mean and two standard deviations across 100 simulations, raw data per simulation is shown by individual points. **B**. In another series of simulations, we varied the effect size of the alleles segregating in the simulated *X. malinche* population from 3-10% of the QTL effect size (red). Simulations with no segregating variation in *X. malinche* are shown in blue for comparison. Large points and whiskers show mean and two standard deviations across 100 simulations, raw data per simulation is shown by individual points. See Supporting Information 10 for a complete description of these simulations.

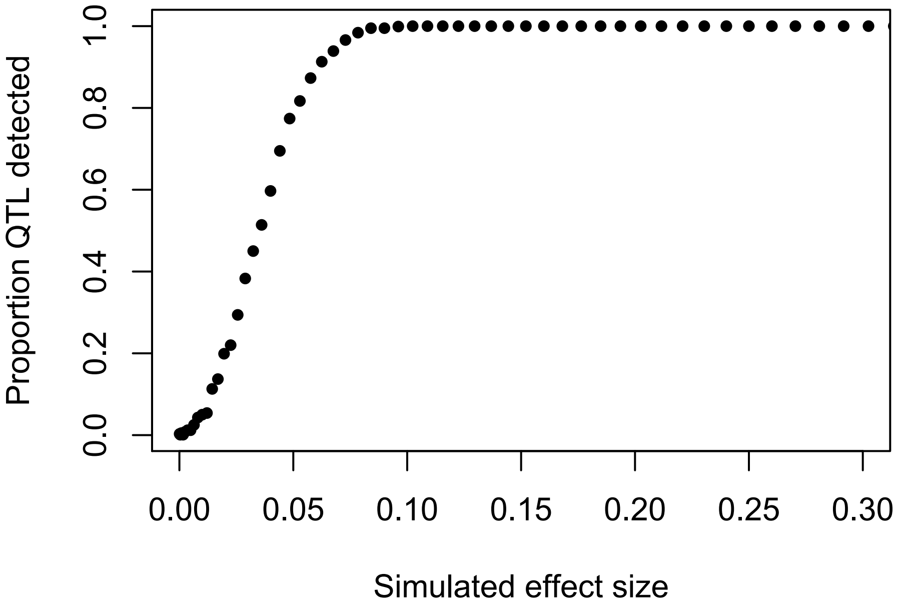


**Fig S16. Predicted power curve for QTL detection in our study as a function of simulated effect size.** We varied the proportion of phenotypic variation explained by a single QTL and tested our power to detect it in simulations. Each point represents the proportion of 1,000 simulations in which the QTL was detected at our genome-wide significance threshold.

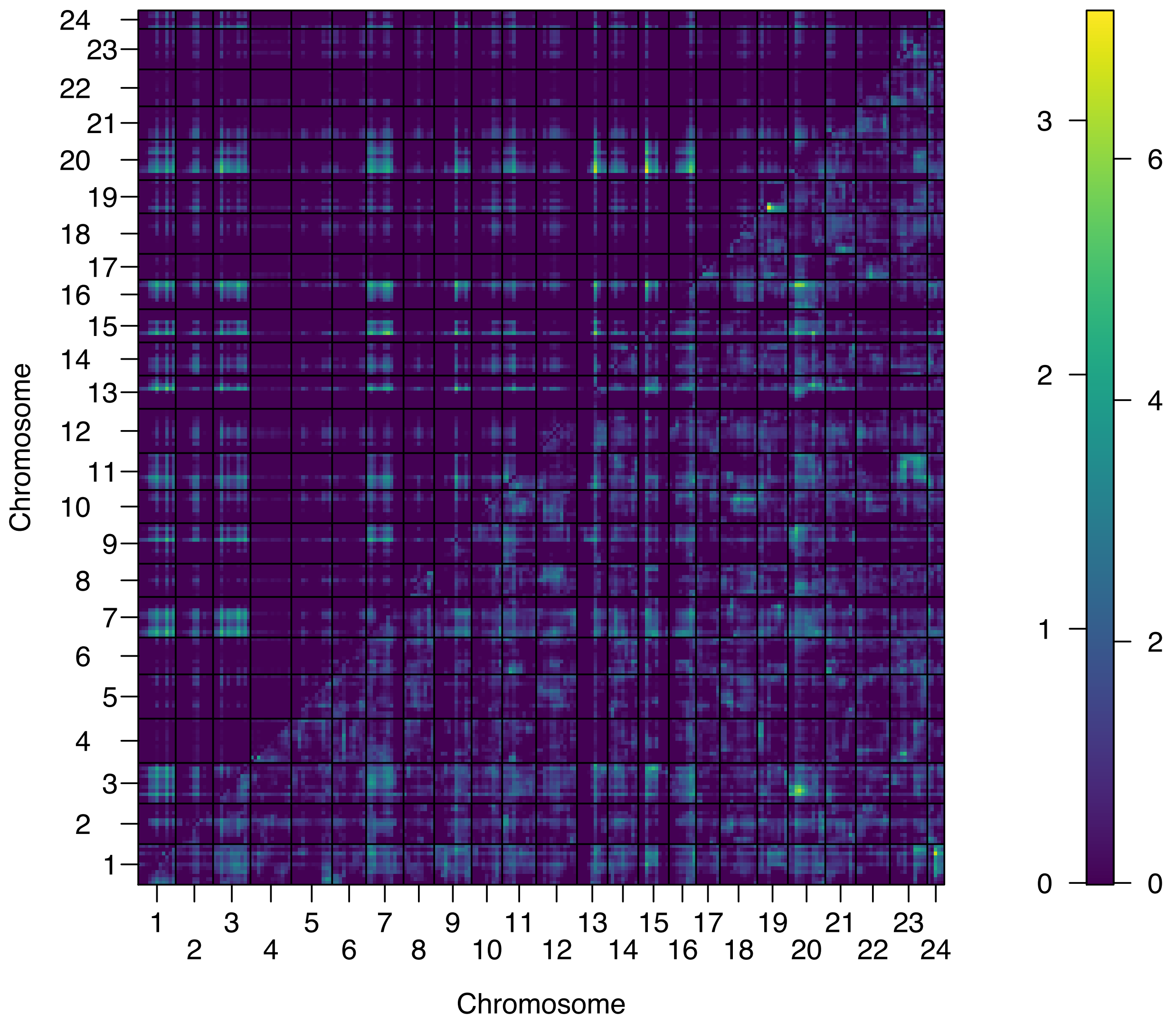


**Fig S17. Results of a two-dimensional two-QTL genome scan for sword length.**
Data shown above and left of the diagonal compare a full two QTL model allowing for epistatic interactions to a single QTL model ($LOD_{fv1}$). Results shown below and right of the diagonal show comparison of a two QTL model, where two single loci on separate chromosomes have additive effects on the phenotype, to a single QTL model ($LOD_{av1}$). The color indicates the magnitude of the LOD score. Numbers on the color scale to the right correspond to values of $LOD_{fv1}$ and $LOD_{av1}$, respectively.

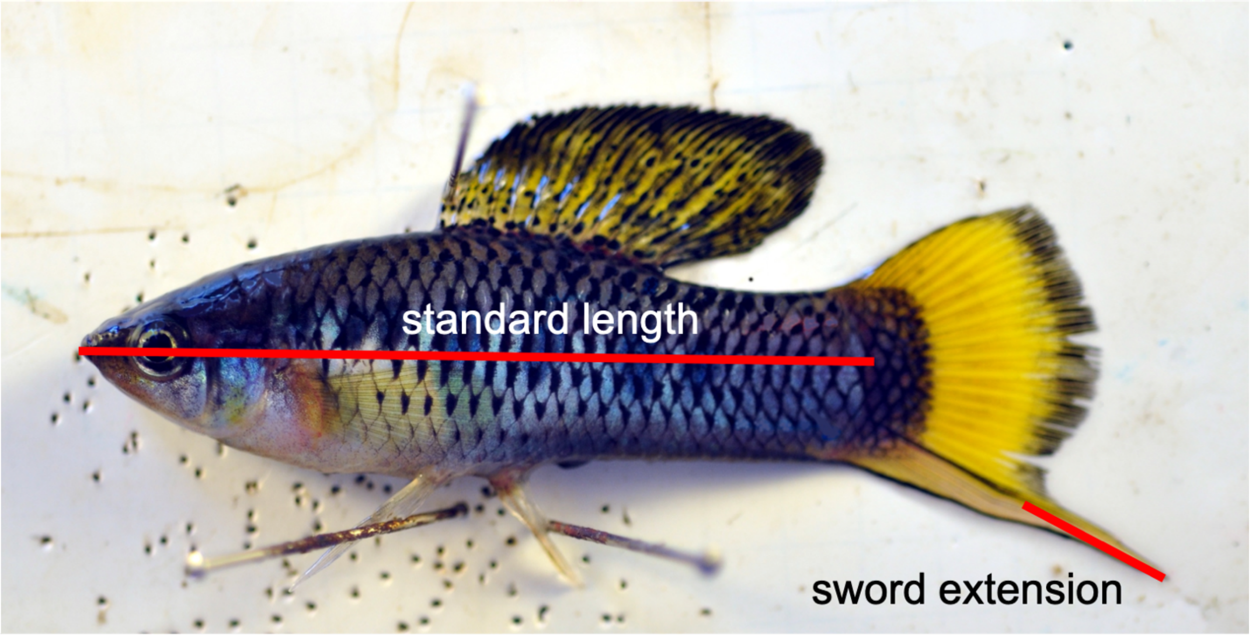


**Fig S18. Schematic showing measurement of sword length using ImageJ software**. Sword extension was normalized by standard length for QTL analysis of sword length.

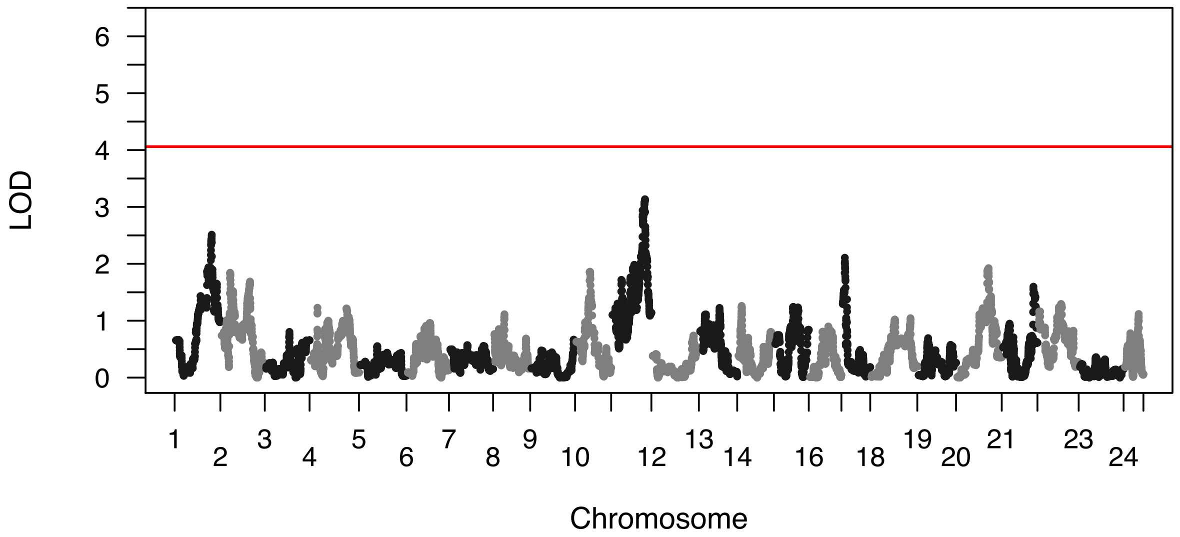


**Fig S19. Manhattan plot showing the results of R/qtl analysis for the presence of the lower sword pigmented edge.** Sword length is one of several phenotypes that makes up the composite sword trait (Fig. S4). The upper sword edge and lower sword edge are also important components of the trait. No genome-wide significant QTL were identified for the lower sword edge trait.

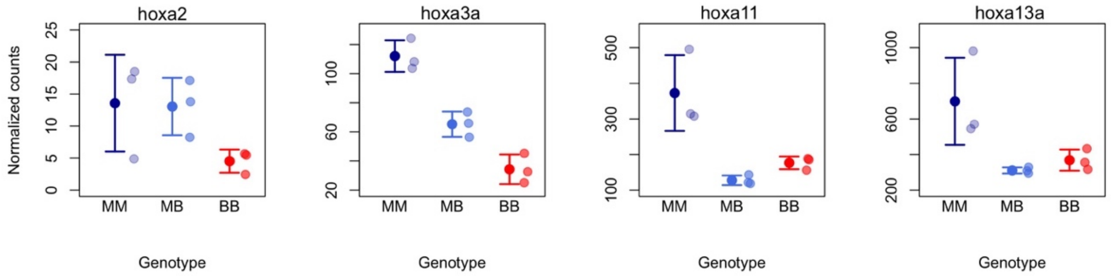


**Fig S20. Expression levels of differentially expressed *hoxa* genes in regenerating caudal fin tissue in *X. malinche* (dark blue), $F_1$ hybrids (light blue) and *X. birchmanni* (red).**
This cluster of genes is found nearbly the QTL region we identify on chromosome 13. Solid points and whiskers show mean and two standard errors of the mean, semi-transparent points show the raw data per individual.

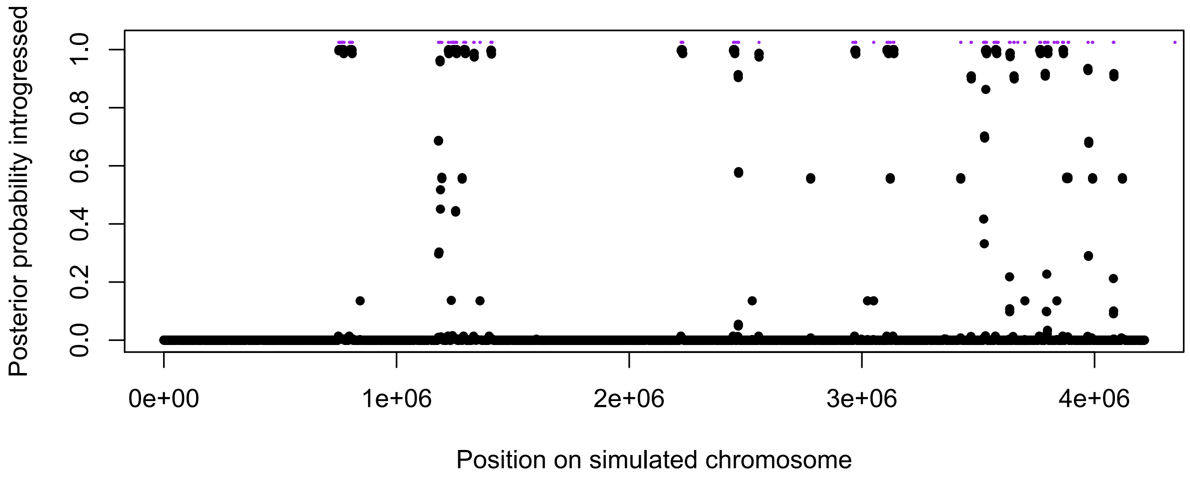


**Fig S21**. **Example of results from simulations of ancient admixture and application of PhyloNet-HMM to infer local ancestry.** Black dots show the posterior probability inferred by PhyloNet-HMM that the site is hybridization derived. Purple lines above the black dots show the true locations of admixture derived tracts, determined from decoding tree sequences in SLiM.

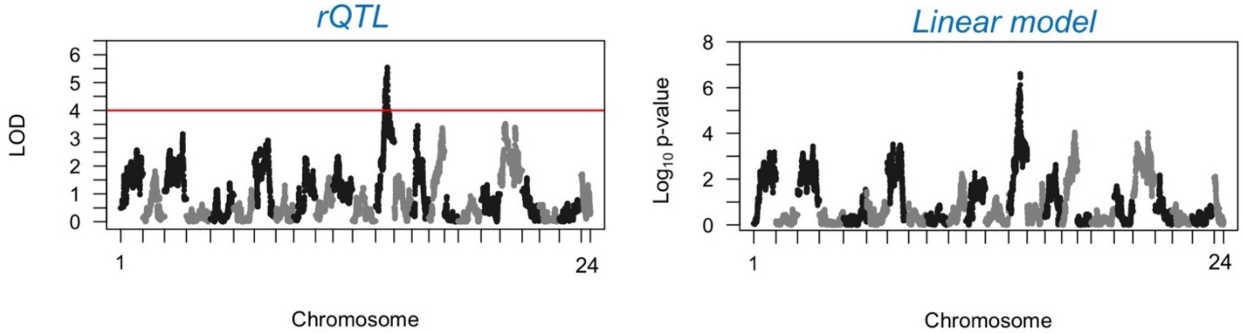


**Fig S22. Comparison of linear model and R/qtl results.** Linear model based mapping results (right) for sword length mirror results from R/qtl (left). Due to extremely long runtimes it was impractical to use R/qtl in simulations but these results suggest that a linear model based approach gives qualitatively similar results.

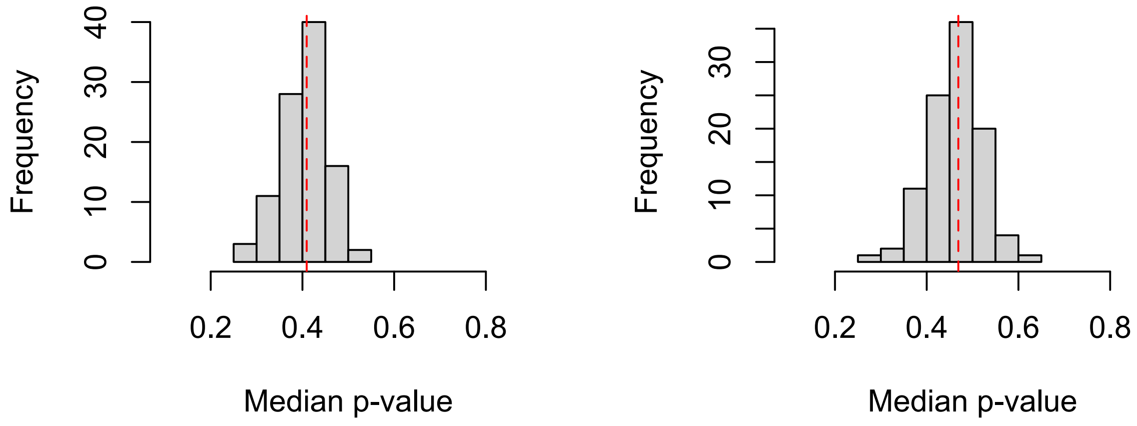


**Fig S23. Distribution of median p-values in simulations of polygenic traits determined by ancestry at 50 underlying loci. A**. Distribution of median p-values at ancestry informative sites in each of 100 simulations when genome-wide ancestry is not included as a covariate in the analysis. **B**. Distribution of median p-values at ancestry informative sites in each of 100 simulations when genome-wide ancestry is included as a covariate in the analysis. Red dashed line indicates the median of each distribution. P-values are skewed towards lower values in **A**. This may reflect lower power when genome-wide ancestry is included as a covariate, or reflect the impacts of uncorrected ancestry structure on p-value distributions.

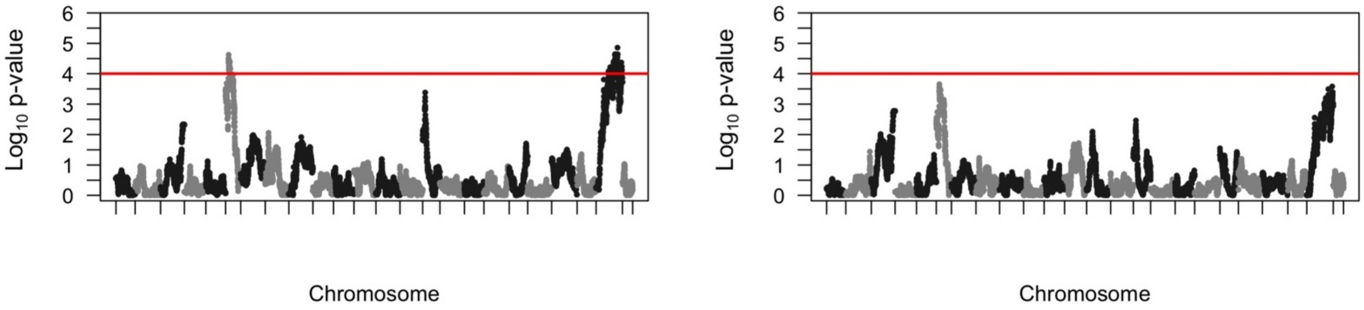


**Fig S24. Example results for simulations of 50 loci contributing to variation in a polygenic trait. A**. Manhattan plot showing results without genome-wide ancestry included as a covariate. **B**. Manhattan plot showing results for the same simulation with genome-wide ancestry included as a covariate. Because we should not have power to detect individual QTL in these simulations, the results in **A** reflect possible inflation of associations when genome-wide ancestry is not accounted for. Red line shows genome-wide significance threshold used in our study.

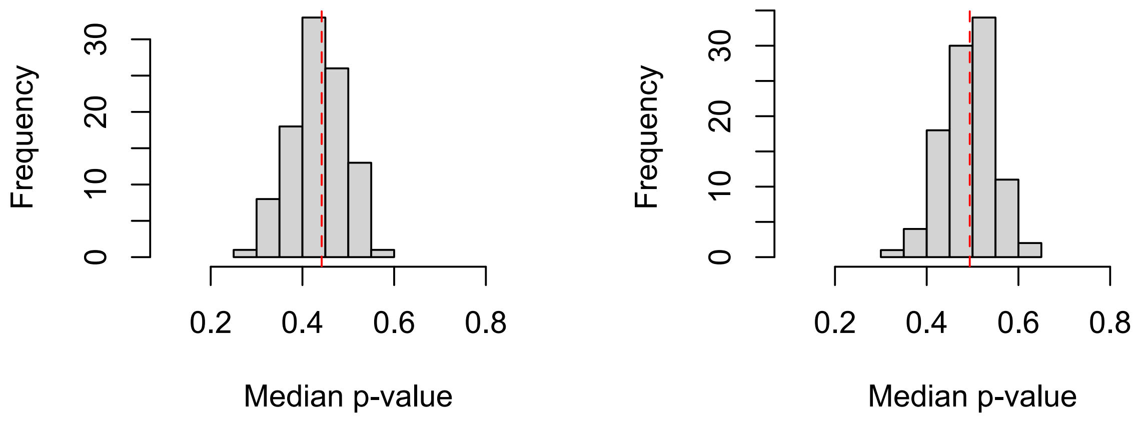


**Fig S25. Distribution of median p-values in simulations of polygenic traits determined by ancestry at 500 underlying loci. A**. Distribution of median p-values at ancestry informative sites in each of 100 simulations when genome-wide ancestry is not included as a covariate in the analysis. **B**. Distribution of median p-values at ancestry informative sites in each of 100 simulations when genome-wide ancestry is included as a covariate in the analysis. Red dashed line indicates the median of each distribution. P-values are significantly shifted towards lower values in **A**. Given that we should have near zero power to detect loci of these effect sizes in our simulations, this skew likely reflects the impacts of uncorrected ancestry structure on p-value distributions.

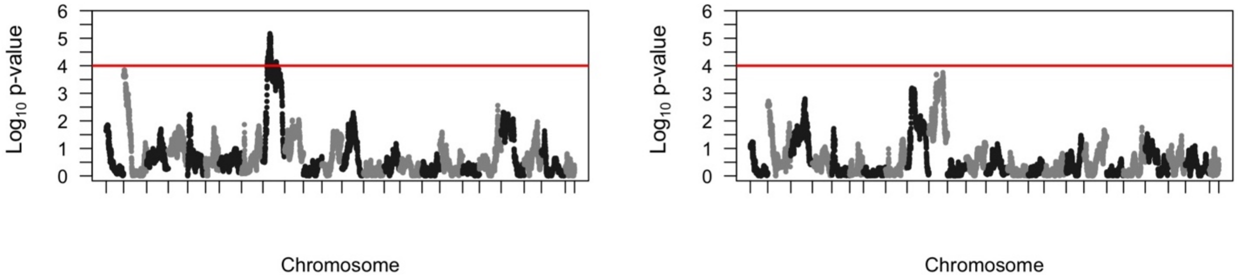


**Fig S26. Example results for simulations of 500 loci contributing to variation in a polygenic trait. A**. Manhattan plot showing results without genome-wide ancestry included as a covariate. **B**. Manhattan plot for the same simulation showing results with genome-wide ancestry included as a covariate. Because we should not have power to detect individual QTL in these simulations, the results in **A** reflect possible inflation of associations when genome-wide ancestry is not accounted for. Red line shows genome-wide significance threshold used in our study.

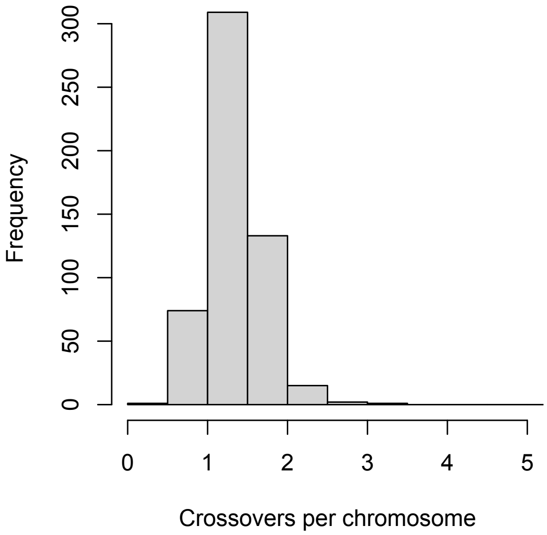


**Fig S27. Distribution of average crossover number across 24 chromosomes in the lab-generated hybrids used in this study.** The mean number of crossovers per chromosome in this dataset was 1.3, similar to observed values in $F_2$ hybrids (1.2), suggesting that the majority of individuals included in our mapping population were $F_2$ hybrids.

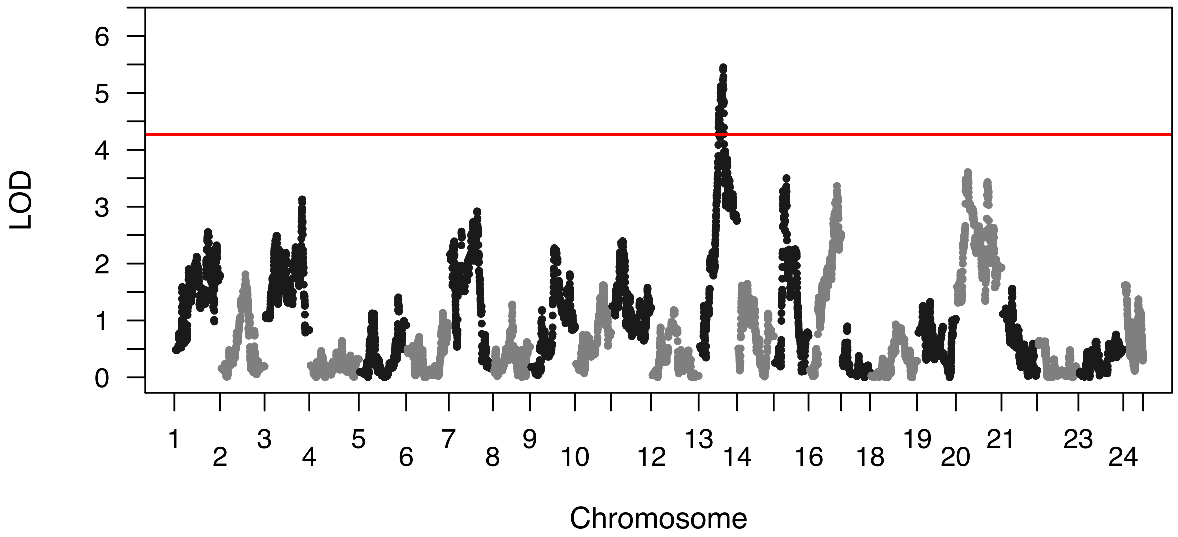


**Fig S28. Manhattan plot of R/qtl mapping results for sword length excluding $F_3$ or later individuals.** Individuals that are likely $F_3$ hybrids given the number of observed crossover events (Fig S27) were excluded from this analysis. Red line indicates genome-wide significant threshold determined by permutation.

### Supplementary Tables

**Tables S1-S3 and S9 are provided as attached excel documents**

**Table S1**. **Genes differentially expressed in regenerated sword tissue.** Information on 3,333 significantly differentially expressed genes between *X. birchmanni* and *X. malinche* in the regenerating caudal fin.

**Table S2**. **All genes that overlap with the joint QTL region**. Genes that fall within the narrowest joint QTL interval on chromosome 13.

**Table S3**. **Summary of expression, annotation, and substitution evidence for candidate genes in the QTL interval.** Evidence associated with the strongest candidates within the QTL region on chromosome 13, those associated with fin or limb phenotypes, growth, skeletal or muscle phenotypes.

**Table S9**. **Overrepresented Gene Ontology terms in the significantly differentially expressed genes between *X. malinche* vs *X. birchmanni* regenerating fin tissue.** Of 3368 GO terms tested, 216 terms were found to be significantly overrepresented with a p-value < 0.05. This analysis included all genes at FDR adjusted p-value < 0.1.

**Table S4. Summary of evolutionary analyses for top candidates.** SIFT, joint ancestry, and dN/dS analysis at other candidate genes identified in the chromosome 13 QTL region. Joint ancestry analysis was conducted using four hybrid populations and comparing observed *X. malinche* ancestry across populations to that expected from randomly sampling windows for each population.

| **Candidate gene** | **Joint ancestry simulation p-value** | **dN/dS** | **SIFT derived in *birchmanni*** |
|---|---|---|---|
| sp4 | 0.04 | NA (dS=0) | One substitution predicted tolerated<br><br>One substitution predicted not tolerated: G710S |
| **sp8** | **0.002** | **0.78** | **Three substitutions predicted tolerated**<br><br>**Two substitutions predicted not tolerated: Y109F, T225I** |
| twistnb | 0.007 | 0.48 | No derived substitutions |
| btd | 0.08 | 3.76 | Two substitutions predicted tolerated |
| grn | 0.095 | 0 | Not applicable |
| cdk6 | 0.02 | 0.53 | One substitution predicted tolerated |
| calcr | 0.004 | 0.064 | One substitution predicted tolerated |
| col1a2 | 0.004 | 0.60 | Two substitutions predicted tolerated |
| dync1i1 | 0.71 | 0.12 | One substitution predicted not tolerated: V526I |

**Table S5**. **Summary of ancestry change analyses for top candidates.** Ancestry changes at other candidate genes identified in the chromosome 13 QTL interval over time in Acuapa time series data.

| **Gene** | **Start position** | **Stop position** | **Change in *X. malinche* ancestry (Acuapa)** |
|---|---|---|---|
| sp4 | 15692146 | 15703983 | -0.17 |
| sp8 | 15715073 | 15716620 | -0.17 |
| twistnb | 15774748 | 15777863 | -0.17 |
| btd | 15837770 | 15841935 | -0.10 |
| grn | 15874280 | 15877603 | 0.01 |
| cdk6 | 15892602 | 15930545 | -0.07 |
| calcr | 16100007 | 16130977 | -0.02 |
| col1a2 | 16245414 | 16260600 | -0.02 |
| dync1i1 | 16368810 | 16428301 | -0.10 |

**Table S6**. **Read information for RNAseq analysis.** Number of reads collected per individual included in RNAseq-based analysis of sword regeneration.

| Sample ID | Group | Day | Total paired-end reads |
|---|---|---|---|
| Xm.1 | *X. malinche* | 10 | 14,110,878 |
| Xm.2 | *X. malinche* | 10 | 12,548,916 |
| Xm.3 | *X. malinche* | 10 | 14,685,792 |
| Xbir.1 | *X. birchmanni* | 10 | 16,798,656 |
| Xbir.2 | *X. birchmanni* | 10 | 13,837,382 |
| Xbir.3 | *X. birchmanni* | 10 | 13,991,052 |
| F1.1 | F1 hybrid | 10 | 10,880,042 |
| F1.2 | F1 hybrid | 10 | 12,845,918 |
| F1.3 | F1 hybrid | 10 | 15,998,254 |

**Table S7. SRA accessions for previously published datasets used in phylogenetic analysis.** Average per basepair coverage when mapped to the *X. birchmanni* reference genome is listed.

| Species | SRA accession | Coverage |
|---|---|---|
| *X. hellerii* | SRR7532852 | 85 |
| *X. maculatus* | SRR7532852 | 91 |
| *X. montezumae* | SRR3086791 | 21 |
| *X. nezahualcoyotl* | SRR3086878 | 24 |
| *X. malinche* | SRR6649369 | 36 |

**Table S8**. **KEGG pathway enrichment in significantly differentially expressed genes between *X. malinche* vs *X. birchmanni* regenerating fin tissue.** Only two pathways were enriched for higher expression in *X. birchmanni*, suggesting that either *X. birchmanni* upregulates or *X. malinche* downregulates (or has constitutively lower expression) of genes involved in ECM-receptor interaction and focal adhesion. This analysis included all genes at FDR adjusted p-value < 0.1.

| | **Mean difference between *Xmal* and *Xbir*** | **p.val** | **q.val** | **set.size** |
|---|---|---|---|---|
| ECM-receptor interaction (xma04512) | -4.3 | $3^{-5}$ | 0.002 | 36 |
| Focal adhesion (xma04510) | -3.6 | 0.0002 | 0.009 | 64 |

### Supporting Information References


1. Schumer M, Powell DL, Corbett-Detig R. Versatile simulations of admixture and accurate local ancestry inference with mixnmatch and ancestryinfer. bioRxiv. 2019; 860924. doi:10.1101/860924

2. Corbett-Detig R, Nielsen R. A Hidden Markov Model Approach for Simultaneously Estimating Local Ancestry and Admixture Time Using Next Generation Sequence Data in Samples of Arbitrary Ploidy. PLOS Genetics. 2017;13: e1006529. doi:10.1371/journal.pgen.1006529

3. Amores A, Catchen J, Nanda I, Warren W, Walter R, Schartl M, et al. A RAD-Tag Genetic Map for the Platyfish (Xiphophorus maculatus) Reveals Mechanisms of Karyotype Evolution Among Teleost Fish. Genetics. 2014;197: 625–641. doi:10.1534/genetics.114.164293

4. Schartl M, Walter RB, Shen Y, Garcia T, Catchen J, Amores A, et al. The genome of the platyfish, Xiphophorus maculatus, provides insights into evolutionary adaptation and several complex traits. Nature Genetics. 2013;45: 567.

5. Schartl M, Kneitz S, Ormanns J, Schmidt C, Anderson, JL, Amores, A, et al. The developmental and genetic architecture of the sexually selected male ornament of swordtails. bioRxiv. 2020.

6. Stelzer G, Rosen N, Plaschkes I, Zimmerman S, Twik M, Fishilevich S, et al. The GeneCards Suite: From Gene Data Mining to Disease Genome Sequence Analyses. Current Protocols in Bioinformatics. 2016;54: 1.30.1-1.30.33. doi:10.1002/cpbi.5

7. Xu S. Theoretical Basis of the Beavis Effect. Genetics. 2003;165: 2259–2268.

8. Otto SP, Jones CD. Detecting the Undetected: Estimating the Total Number of Loci Underlying a Quantitative Trait. Genetics. 2000;156: 2093–2107.

9. Broman KW, Wu H, Sen Ś, Churchill GA. R/qtl: QTL mapping in experimental crosses. Bioinformatics. 2003;19: 889–890. doi:10.1093/bioinformatics/btg112

10. Bray NL, Pimentel H, Melsted P, Pachter L. Near-optimal probabilistic RNA-seq quantification. Nat Biotechnol. 2016;34: 525–527. doi:10.1038/nbt.3519

11. Chiaromonte F, Miller W, Eric E. Bouhassira. Gene Length and Proximity to Neighbors Affect Genome-Wide Expression Levels. Genome Res. 2003;13: 2602–2608. doi:10.1101/gr.1169203

12. Szklarczyk D, Morris JH, Cook H, Kuhn M, Wyder S, Simonovic M, et al. The STRING database in 2017: quality-controlled protein–protein association networks, made broadly accessible. Nucleic Acids Res. 2017;45: D362–D368. doi:10.1093/nar/gkw937

13. van de Geijn B, McVicker G, Gilad Y, Pritchard JK. WASP: allele-specific software for robust molecular quantitative trait locus discovery. Nat Methods. 2015;12: 1061–1063. doi:10.1038/nmeth.3582

14. Walling CA, Royle NJ, Metcalfe NB, Lindström J. Green swordtails alter their age at maturation in response to the population level of male ornamentation. Biol Lett. 2007;3: 144–146. doi:10.1098/rsbl.2006.0608

15. Schumer M, Xu C, Powell DL, Durvasula A, Skov L, Holland C, et al. Natural selection interacts with recombination to shape the evolution of hybrid genomes. Science. 2018;360: 656. doi:10.1126/science.aar3684

16. Haller BC, Messer PW. SLiM 3: Forward Genetic Simulations Beyond the Wright–Fisher Model. Hernandez R, editor. Molecular Biology and Evolution. 2019;36: 632–637. doi:10.1093/molbev/msy228

17. Haller BC, Galloway J, Kelleher J, Messer PW, Ralph PL. Tree-sequence recording in SLiM opens new horizons for forward-time simulation of whole genomes. bioRxiv. 2018; 407783. doi:10.1101/407783

18. Liu KJ, Dai J, Truong K, Song Y, Kohn MH, Nakhleh L. An HMM-Based Comparative Genomic Framework for Detecting Introgression in Eukaryotes. PLOS Computational Biology. 2014;10: e1003649. doi:10.1371/journal.pcbi.1003649

19. Patterson N, Moorjani P, Luo Y, Mallick S, Rohland N, Zhan Y, et al. Ancient Admixture in Human History. Genetics. 2012;192: 1065–1093. doi:10.1534/genetics.112.145037

20. Visscher PM, Haley CS. Detection of putative quantitative trait loci in line crosses under infinitesimal genetic models. Theor Appl Genet. 1996;93: 691–702. doi:10.1007/BF00224064

21. Cui R, Schumer M, Rosenthal GG. Admix'em: a flexible framework for forward-time simulations of hybrid populations with selection and mate choice. Bioinformatics. 2016;32: 1103–1105. doi:10.1093/bioinformatics/btv700
